# Molecular analysis of archival diagnostic prostate cancer biopsies identifies genomic similarities in cases with progression post-radiotherapy, and those with *de novo* metastatic disease

**DOI:** 10.1101/2023.09.04.555868

**Authors:** P Charlton, D O’Reilly, Y Philippou, SR Rao, AD Lamb, IG Mills, G Higgins, FC Hamdy, C Verrill, FM Buffa, RJ Bryant

**Author notes:** Corresponding senior authors. Contacts and. Senior authors (FM Buffa, RJ Bryant).

## Abstract

**Purpose:** It is important to identify molecular features that improve prostate cancer (PCa) risk stratification before radical treatment with curative intent. Molecular analysis of historical diagnostic formalin-fixed paraffin-embedded (FFPE) prostate biopsies from cohorts with post-radiotherapy (RT) long-term clinical follow-up has been limited. Utilizing parallel sequencing modalities, we performed a proof-of-principle sequencing analysis of historical diagnostic FFPE prostate biopsies. We compared patients with i) stable PCa post-primary or salvage RT (sPCa), ii) progressing PCa post-RT (pPCa), and iii) *de novo* metastatic PCa (mPCa).

**Experimental Design:** A cohort of 19 patients with diagnostic prostate biopsies (n=6 sPCa, n=5 pPCa, n=8 mPCa) and mean 4 years 10 months follow-up (diagnosed 2009-2016) underwent nucleic acid extraction from demarcated malignancy. Samples underwent 3’RNA sequencing (3’RNAseq) (n=19), nanoString analysis (n=12) and Illumina 850k methylation (n=8) sequencing. Bioinformatic analysis was performed to coherently identify differentially expressed genes (DEGs) and methylated genomic regions (MGRs).

**Results:** 18 of 19 samples provided useable 3’RNAseq data. Principal Component Analysis (PCA) demonstrated similar expression profiles between pPCa and mPCa cases, versus sPCa. Coherently differentially methylated probes between these groups identified ∼600 differentially MGRs. The top 50 genes with increased expression in pPCa patients were associated with reduced progression-free survival post-RT (*p*<0.0001) in an external cohort.

**Conclusions:** 3’RNAseq, nanoString and 850K-methylation analyses are each achievable from historical FFPE diagnostic pre-treatment prostate biopsies, unlocking the potential to utilize large cohorts of historic clinical samples. Profiling similarities between individuals with pPCa and mPCa suggests biological similarities and historical radiological staging limitations, which warrant further investigation.

## Introduction

Localized PCa may be treated with curative intent by radical surgery, or RT with concomitant androgen-deprivation therapy (ADT), with equivalent rates of post-treatment disease progression, but differences in side effect profiles, at 15-years clinical follow-up^1,2^. In the clinical oncology setting, it is recognized that at long-term follow-up a group of patients will experience disease progression following RT^3^. In addition to use of clinical risk scores based on PSA, tumor grade and stage^3,4^, baseline molecular characterisation of diagnostic biopsies offers the potential to identify patients at high risk of post-RT relapse. This approach may facilitate more accurate risk-stratification in the immediate post-diagnosis pre-treatment space, to enable more appropriate treatment selection.

Several prognostic and predictive transcriptomic classifiers have been developed for PCa. However, none are routinely used in the clinic in the pre-RT setting. Currently available classifiers include the Oncotype, Decipher, Prolaris, metastatic and hypoxia signatures^5-10^. The Decipher, Oncotype and cell cycle progression (CCP) signatures have demonstrated clinical utility in predicting disease recurrence after both radical surgery and radical RT^8,11-15^, despite these classifiers being derived from surgically treated patients. A prostate-specific molecular signature of hypoxia has been demonstrated to predict biochemical recurrence in the salvage RT setting for local disease recurrence post-radical prostatectomy and is an independent prognostic indicator for patients with localized PCa receiving RT^10^. Methylation-based molecular features of PCa have also been found to be associated with clinical outcome after salvage RT^15-17^.

Use of archival FFPE diagnostic prostate biopsy samples for RNA sequencing is technically challenging given the small volume of available tissue, and the degradation and cross-linking of RNA over time. 3’RNAseq is potentially well suited for sequencing of degraded RNA, utilizing only the 3’ ends of RNA fragments, and providing a single read per gene transcript. NanoString analysis provides an alternative strategy for molecular analysis of clinical samples, as this technique uses reporter probes to hybridize mRNA, and reports fewer genes (∼600 for nanoString, versus ∼ 20,000 for 3’RNAseq), thus significantly reducing computational resources and analysis time. Illumina 850k methylation analysis has greater power over previous methylation arrays due to the increased number of probes. Utilizing these distinct molecular analysis technologies in an orthogonal manner may identify biologically relevant genes, versus each single genomic technique in isolation.

Previous studies in the field have analyzed the molecular features of clinical samples from patients with localized PCa treated with RT, to derive a signature associated with clinical outcome, such as those described as being hypoxic or metastatic^9,10^. In this study we investigated whether molecular features associated with subsequent development of metastatic disease could be identified in the historical diagnostic FFPE prostate biopsy samples from patients with long-term clinical follow-up after primary radical or salvage RT. We used orthogonal analyses of a carefully curated small number of samples and included cases with baseline *de novo* mPCa at presentation as a comparator. Using this approach, we aimed to provide proof-of-concept that this technique could unlock the door to future larger scale studies from mature cohorts, such as those from the ProtecT (Prostate testing for cancer and Treatment) study^1^, with fifteen years clinical follow-up. Such studies have the potential to provide added-value in future risk-stratification for men newly diagnosed with intermediate- or high-risk localized or locally advanced PCa undergoing radical radiotherapy, based on a personalized medicine genomic analysis of baseline molecular features of each malignancy, alongside conventional risk parameters such as PSA, tumour grade and stage^3,18-20^.

## Methods

### Patient Identification

Baseline diagnostic prostate biopsy samples from patients in the ProMPT (Prostate cancer: Mechanisms of Progression and Treatment) cohort with carefully curated clinicopathological features and long-term clinical follow-up following primary or salvage RT were identified. To investigate baseline molecular features of development of mPCa after RT, three groups of patients were identified for this proof-of-concept study: individuals with i) sPCa, ii) pPCa, and iii) mPCa (**Figure 1** and Figure S1).

**Figure 1.**
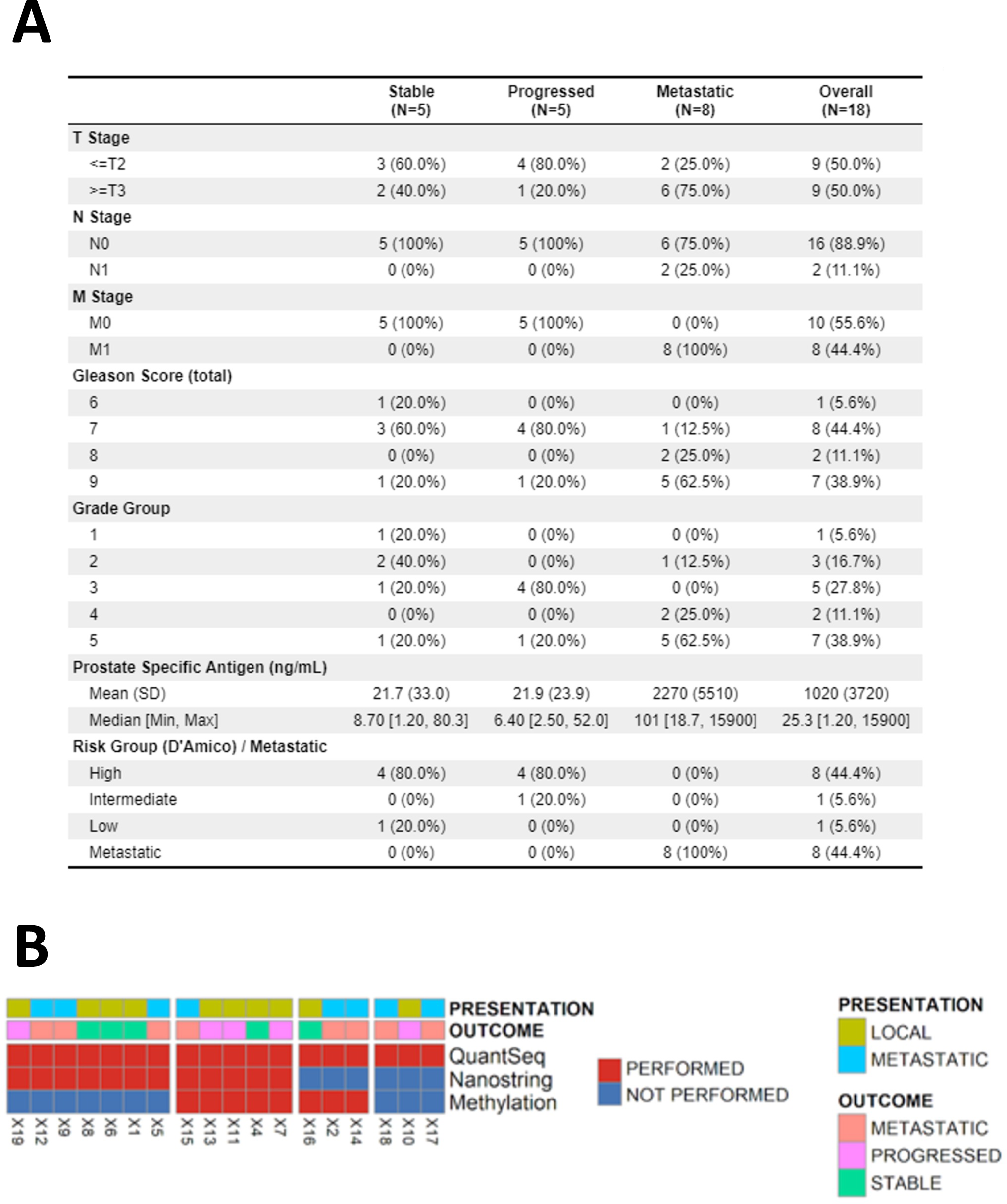
Summary of clinical characteristics of the cohort. (**A**). Heatmap summary of the sequencing performed on the cohort (**B**). Quantseq 3’RNA sequencing was performed for all cases; nanoString analysis was performed for 12 of the cases; DNA methylation analysis was performed for 8 of the cases.

### Pathology & Nucleic acid extraction

Pathology sections were reviewed by a specialist uropathologist for tissue availability, Gleason Grade Group, and percentage tumor content. PCa was annotated in prostate biopsy samples on a single slide, and four 5 µm serial sections per patient were macro-dissected using a sterile scalpel. Samples were pooled per patient, and RNA extracted using the Roche HighPure FFPE kit. 8 patient samples contained sufficient material for 4 further slides to be utilised for DNA extraction using the ROCHE FPPE high-pure DNA kit protocol.

### Sequencing techniques

3’RNAseq sequencing library preparation was performed with QuantSeq 3’-mRNA-Seq Library Prep Kit (Lexogen), and libraries were sequenced on Illumina NextSeq flow cells (75 bp fragments) at the Welcome Trust Centre for Human Genetics (University of Oxford, UK). For nanoString analysis, 12 samples with optimal spectrophotometric characteristics (determined by NanoDrop) were selected (from 4 individuals with each of sPCa, pPCa, and mPCa). Samples underwent nCounter® Human PanCancer Panel Gene Expression profiling according to the manufacturer’s instructions, and data was acquired using the nCounter® SPRINT profiler. For DNA methylation analysis, bisulphite conversion was performed using the EZ-96 DNA Bisulphite Zymo Research conversion protocol. The Illumina Infinium HD Methylation Assay protocol was followed, and samples were hybridized to Human Methylation EPIC beadchips (Genomics Birmingham, UK).

### Gene Expression & Methylation Data Analysis

3’RNAseq FastQ files were concatenated for each patient, and polyA and llumina adapter trimming (AGATCGGAAGAGC) was performed using trimmomatic (v0.25), prior to alignment with STAR (2.7.7a) to Hg38 with featurecounts used to generate counts (**Supplementary Methods**)^21^. Filtering of lowly expressed genes was performed, followed by differential expression analysis using DESEQ2 (v1.26.0)^22^. Over-representation analysis was performed using Clusterprofiler, enrichr and gprofiler. The gene set variation analysis GSVA (v1.34.0) package was used to perform single sample gene set enrichment analysis (ssGSEA)^23^.

### Quality Control

3’RNAseq quality control (QC) was performed with FastQC pre- and post-trimming. One sample failed sequencing, with <100,000 counts, and was not used for further analysis. Although not strictly required for DESeq2 analysis, counts were filtered to only include genes with >1 count in ≥5 samples (minimum group size). For nanoString analysis, data were imported into nSolver^TM^ analysis software v2.5, and QC performed according to nanoString guidelines, with gene transcripts normalized to housekeeping genes. For DNA methylation analysis, QC, filtering of poor performing probes, cross-reactive probes and normalization was performed prior to differential methylation analysis using minfi package (v1.32.0).

As a measure of the performance characteristics of the dataset, the SigQC protocol^24^ was used to generate QC metrics for previously validated signatures (DECIPHER, Prolaris, Oncotype, and prostate hypoxia, each obtained from the published literature^5-10^ and these were compared to two previously derived ‘gold standard’ datasets (the TCGA and Jain RT datasets). Sigcheck (package v2.18.0) was used to assess the performance of these gene sets compared to random genes and known signatures with 1000 iterations^25^. KMunicate and survival packages were used for survival analysis.

### Statistical Survival Analysis

Sequencing and clinical data from the external dataset GSE116918 were downloaded from Gene Expression Omnibus (GEO) repository (https://www.ncbi.nlm.nih.gov/geo/) and survival analysis performed using a Cox proportional hazards model comprising clinicopathological features and expression of selected genes, followed by estimation of time-dependent Receiver Operator Characteristics (ROC) using TimeROC (package 0.4)^26^ (**Supplementary Methods**).

## Results

Samples from 18 of 19 patients were included in this study (n=6 with sPCa, n=5 with pPCa, and n=8 with mPCa). One sample failed 3’RNAseq and was excluded from further analysis. Baseline clinicopathological features of the cohort, and a summary of the sequencing technologies (n=18 3’RNAseq; n=12 nanoString; n=8 Methylation) used per patient, are shown in **Figure 1** and Table S1. Baseline clinicopathological features of the two patient groups receiving RT (i.e. those with sPCa, and those with pPCa) were similar, with a similar mean PSA, and both groups contained n=4 D’Amico high-risk patients.

The sigQC protocol was used for QC to evaluate the performance of previously validated PCa signatures (prostate hypoxia, Decipher, Prolaris and Oncotype) in the sequencing dataset, compared to the “gold standard” TCGA prostate (pan-cancer) dataset, and an external dataset of patients with localized PCa treated with radical radiotherapy by Jain (GSE116918)^11^. QC metrics demonstrated that these signatures perform at a comparable level in the dataset compared to previously published datasets (Figures S1-S3), indicating the general applicability of 3’RNAseq technology, and providing a validation of the utility of the dataset.

PCA of the 3’RNAseq data demonstrated some separation of the sPCa cases from those with pPCa or mPCa (**Figure 2A**). A comparison of sPCa versus pPCa cases identified 558 DEGs (*padj* < 0.05, LFC >1 or <-1) (n=432 increased expression, n=126 decreased expression) (**Figure 2B**). A similar number of DEGs were observed between mPCa and sPCa cases (n=422 increased expression, n=95 decreased expression) (**Figure 2B**). Only one gene was significantly differentially expressed between pPCa and mPCa cases (**Figure 2B**). The majority of DEGs between sPCa versus pPCa, and sPCa versus mPCa, were observed to have concordant directionality (n=421 increased expression, n=92 decreased expression) (**Figure 2C**).

**Figure 2.**
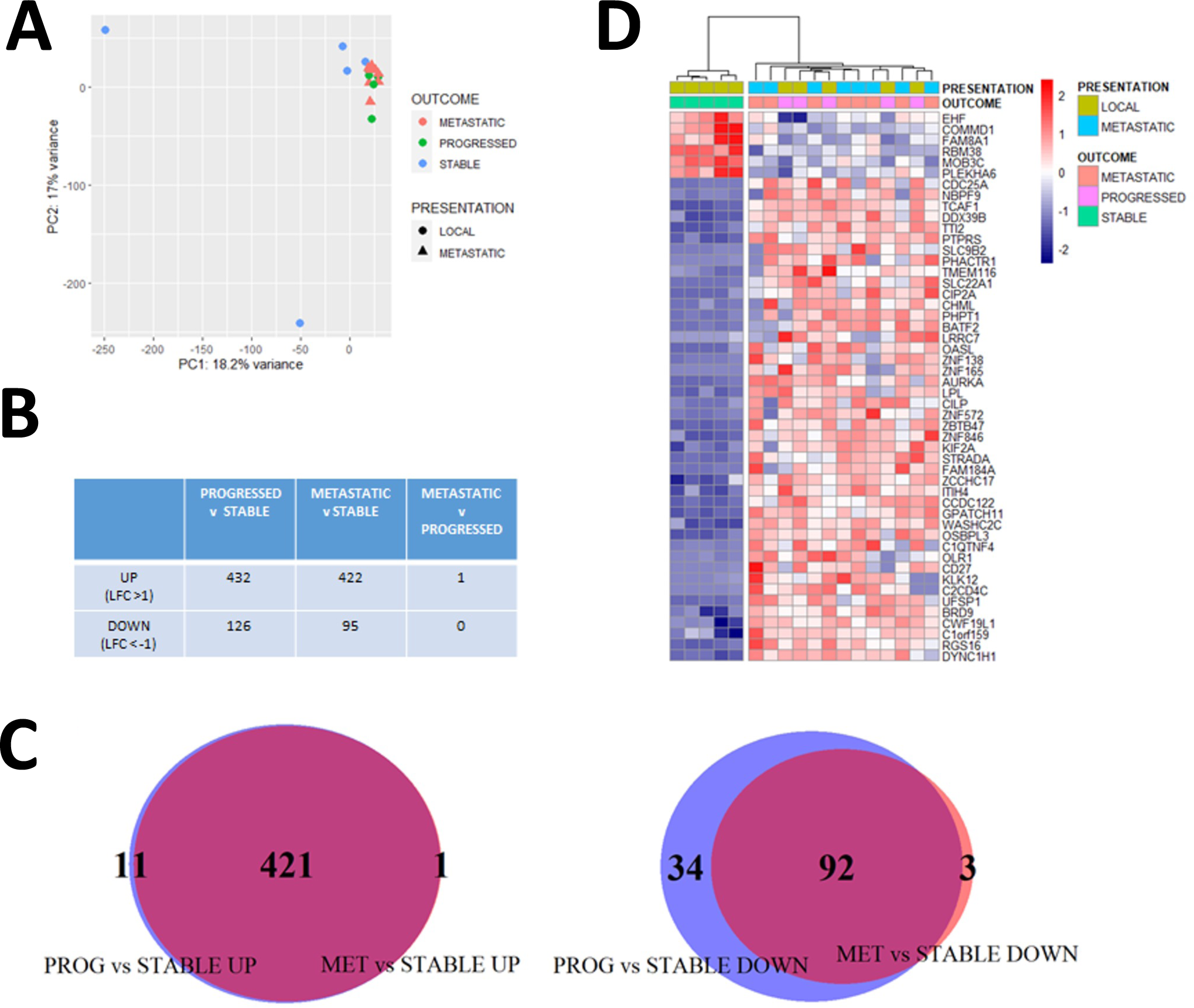
3’RNAseq analysis of the cohort. Principal Component Analysis (PCA) of normalised (rlog) gene expression data demonstrated some separation of stable cases from the progressed and *de novo* metastatic patients. (**A**). The number of differentially expressed genes (DEGs) between comparisons of stable, progressed, and *de novo* metastatic cases is shown in (**B**) (adjusted *p*-value <0.05, logfold change >1/< -1). A Venn diagram Venn Diagram demonstrated the overlap of DEGs in (**B**) by direction, increased (left panel) between progressed versus stable cases (PROG vs STABLE UP) and *de novo* metastatic versus stable cases (MET vs STABLE UP), and decreased (right panel) between progressed versus stable cases (PROG vs STABLE DOWN) and *de novo* metastatic versus stable cases (MET vs STABLE DOWN) (**C**). This analysis identified common DEGs genes between groups (**C**). Heatmap of DEGs in stable versus progressed cases (adjusted *p*-value < 0.05). Hierachical clustering (ward.D2 method and euclidean distance) demonstrated clustering of the progressed and *de novo* metastatic cases, which have a similar gene expression pattern (**D**).

Hierarchical clustering of the top 50 DEGs between sPCa and pPCa cases, demonstrated that pPCa and mPCa samples had a similar expression pattern, and this was distinct from sPCa cases (**Figure 2D**), supporting the PCA findings. Gene ontology over-representation analysis of increased DEGs in sPCa versus pPCa cases (*padj* <0.01), and sPCa versus mPCa (*padj* <0.05) cases, identified pathways associated with spindle pole and centrosome function respectively (Figures S4-S5).

NanoString analysis performed on 12 cases (4 from each of the three groups) demonstrated similar differences between sPCa cases and those with either pPCa or mPCa. PCA demonstrated separation of sPCa from pPCa and mPCa cases (except for one metastatic case) (**Figure 3A**). The top 25 DEGs by NSolver^TM^ analysis between sPCa and pPCa cases (*p*<0.05) (**Figure 3B**) demonstrated a similar expression pattern difference to that observed between sPCa and mPCa samples (**Figure 3C**), however this result was not statistically significant on correction for multiple testing (Table S2). Non-hierarchical clustering of log-normalized nanoString expression data for these genes demonstrated that sPCa cases clustered with one mPCa case (**Figure 3B**). Similar findings were observed in the comparison of sPCa versus mPCa cases (**Figure 3C**). 16 of the top 25 DEGs (64%) were common between the two comparisons (**Figure 3D**).

**Figure 3.**
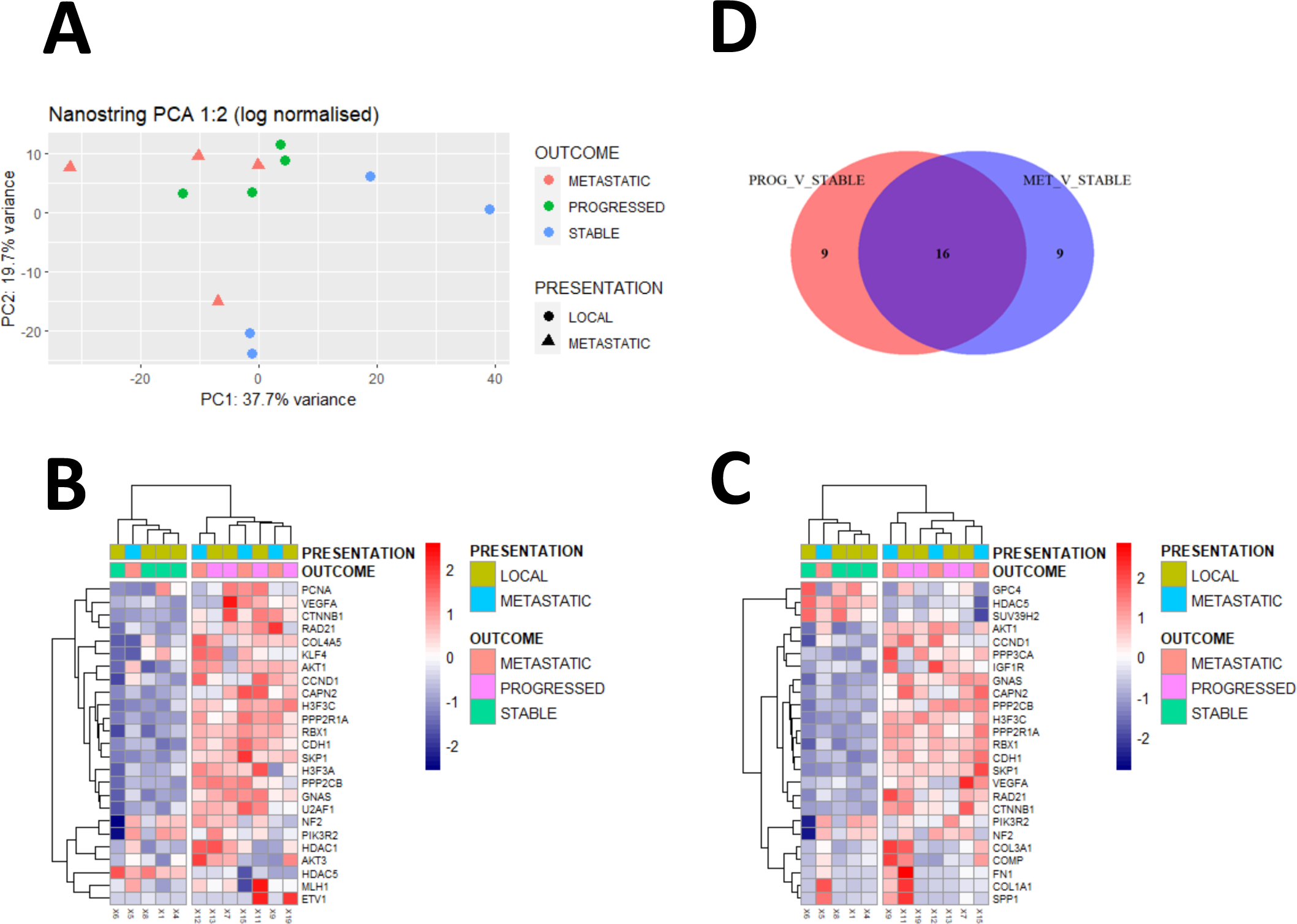
nanoString analysis of the cohort. Principal Component Analysis (PCA, 1:2) of normalised (log) nanoString gene expression data demonstrated some separation of stable cases from the progressed and *de novo* metastatic cases. (**A**). Heatmap of the top 25 DEGs identified by nanoString Nsolver analysis between stable and progressed cases (**p** < 0.05) (**B**). Non-hierarchical clustering (ward.D2 method and euclidean distance) demonstrated clustering of progressed and *de novo* metastatic patients with a similar expression pattern (**B**). A heatmap of the top 25 DEGs (*p* < 0.05) between *de novo* metastatic and stable cases (identified in **B**) demonstrated clustering and similarity of the progressed and *de novo* metastatic cases (**C**). A Venn diagram demonstrated the overlap between the top 25 DEGs between progressed versus stable cases (PROG_V_STABLE), and *de novo* metastatic versus stable cases (MET_V_STABLE) (**D**).

A comparison of nanoString expression (log-normalized) and 3’RNAseq expression (rlog-normalized) yielded an overall Spearman correlation coefficient of 0.68. We identified an overlap of 4 directionally concordant DEGs between sPCa and pPCa cases (increased DEGs: *GNAS*, *ETV1*, *COL2A1*; decreased DEGs: *HDAC5*) in both the 3’RNAseq (utilizing less stringent cutoffs, *padj*<0.2) and nanoString (*p*<0.05) platforms (Table S2A).

ssGSEA of Quantseq data identified DEGs in pathways associated with metastasis, centrosome, and methylation pathways in sPCa cases versus combined pPCa and mPCa cases, with only the centrosome pathways containing overlapping sets of genes (Figures S6- S7, Table S3).

Taking forward the observation that gene expression profiles were similar in pPCa and mPCa cases, ssGSEA was performed using a previously validated subset of metastatic signatures and pathways from the Molecular Signatures Database (MSigDB) website (https://www.gsea-msigdb.org/gsea/msigdb). Differential expression analysis was performed to compare sPCa versus pPCa, which identified 10 statistically significant signatures (*padj*<0.1) as visualized in the heatmap (**Figure 4A**). PCA of ssGSEA scores demonstrated separation of sPCa cases versus pPCa and mPCa cases (**Figure 4B**).

**Figure 4.**
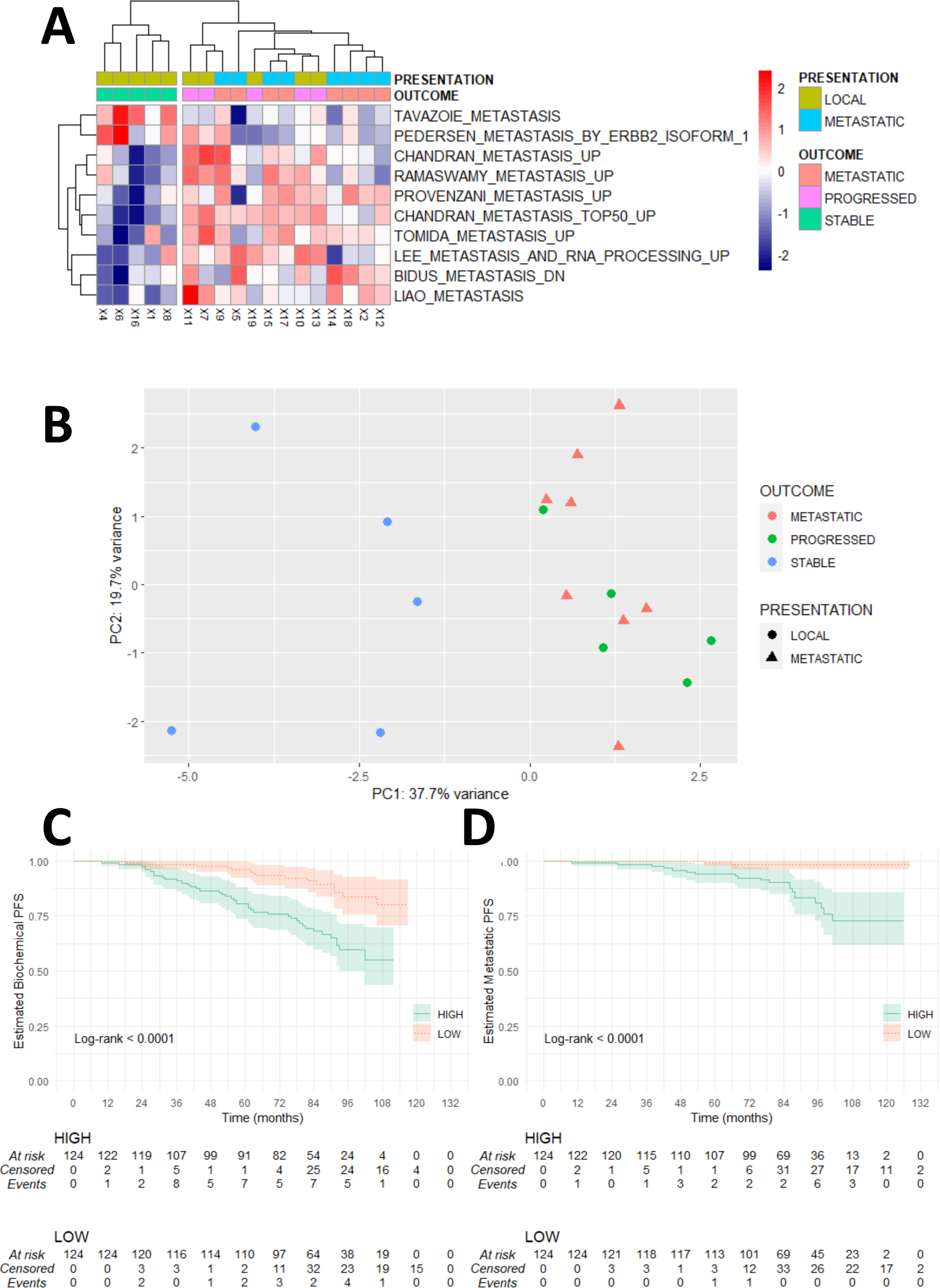
Analysis of findings against a previously validated subset of metastatic signatures and pathways. Guided single sample gene set analysis (ssGSEA) identified differences in metastasis pathways between stable versus progressed and *de novo* metastatic cases (metastatic signatures subset of C2 MsigDB curated pathways, adjusted *p*-value < 0.1) (**A**). Principle component analysis (PCA) of metastasis signature ssGSEA scores defined in **A** demonstrated separation of stable versus progressed and *de novo* metastatic cases (**B**). 36 of the top 50 protein-coding DEGs increased in the progressed versus stable ProMPT cases were then analysed in the external Belfast radiotherapy-treated dataset (GSE116918). This demonstrated an association between the median expression of these DEGs (divided into HIGH vs LOW by median expression of all probes) and biochemical progression-free survival (**C**) and metastatic progression-free survival (**D**).

To explore the potential biological relevance of the DEGs from the 3’RNAseq data, the expression of the top 50 increased protein coding genes in pPCa versus sPCa cases, which were also increased in mPCa versus sPCa cases (**Figure 2D**), was explored in the large external Jain RT dataset^11^, which contains data for 248 PCa patients treated with RT with clinical follow-up. Probes corresponding to the top 50 increased protein-coding genes were selected (37 genes were represented in the dataset by 116 probes, see **Supplementary Methods**), and the cohort was divided into *high* and *low* expression cohorts (using the median as threshold for mean expression of all 37 genes). Biochemical recurrence-free and metastatic progression-free survival curves were observed to be significantly (*p*<0.001) different between the *high* and *low* expression cohorts (**Figures 4C-D**). A significantly increased hazard ratio (HR) in the *high* expression cohort for biochemical (HR 2.5 [1.4-4.3], *p*<0.003) and metastatic (HR 5.1 [1.7-15], p<0.004) progression-free survival was observed on univariable analysis in a cox proportional hazards model (Figures S12 and S13). Signature performance for biochemical recurrence-free survival and metastatic progression-free survival was significant compared to random signatures, cancer signatures, and permutations of survival and feature data performed using the SigCheck package with 1000 iterations (Figures S8-S10). Four genes (*CDC25A*, *OLR1*, *CDON* and *DDX39B*) were found to be independently associated with biochemical progression-free survival (lower limit of Hazard Ratio >1) in a cox proportional hazards model (Figure S12A). A multivariable Cox proportional hazard model sequentially incorporating clinicopathological features using clinically relevant cutoffs (T-stage, Gleason score (dichotomized as sum score 6-7 or sum score 8-10) and PSA (dichotomized as <20, ≥20) and high vs low expression of the 4 genes (mean expression of all 4 genes with the median value used as threshold to divide cohort) demonstrated a hazard ratio of 3.47 [1.79 – 6.7] (p<0.001) for high versus low expression cohorts (Table S4 and Figure S12B). Estimation of time-dependent ROC demonstrated the ability of this 4-gene signature (expressed as mean expression of all 4 genes), with PSA and Gleason scores, as continuous variables in a Cox proportional hazards model to predict for biochemical Area Under the Curve (AUC 76.8) and metastatic (AUC 82.9) progression-free survival (package TimeROC) (Figure S13). The associations of individual genes with clinical characteristics are shown in Table S5.

Methylation analysis was performed on 8 samples (2 with sPCa, 3 with pPCa, and 3 with mPCa) on two illumina 850K EPIC arrays. Following normalization and filtering of methylation data, PCA identified similar findings to those observed in the 3’RNAseq data, with separation of the sPCa cases from pPCa and mPCa cases (**Figure 5A**). Differential methylation analysis (performed with minfi package v1.32.0) identified 1305 probes to be significantly differentially methylated in sPCa versus pPCa cases (*padj*<0.05), and 9551 probes to be significantly differentially methylated in sPCa versus mPCa cases. Most probes (94.5%, 874 of 925 probes) hypomethylated in sPCa versus pPCa cases were also hypomethylated in sPCa versus mPCa cases. Most probes (83.1%, 316 of 380 probes) hypermethylated in sPCa versus pPCa cases were hypermethylated in pPCa versus mPCa cases (**Figure 5B**). This similarity is demonstrated by only 1 probe being significantly differentially methylated between pPCa and mPCa cases and is visualized in heatmaps of the coherently differentially methylated probes (**Figure 5C-5D**).

**Figure 5.**
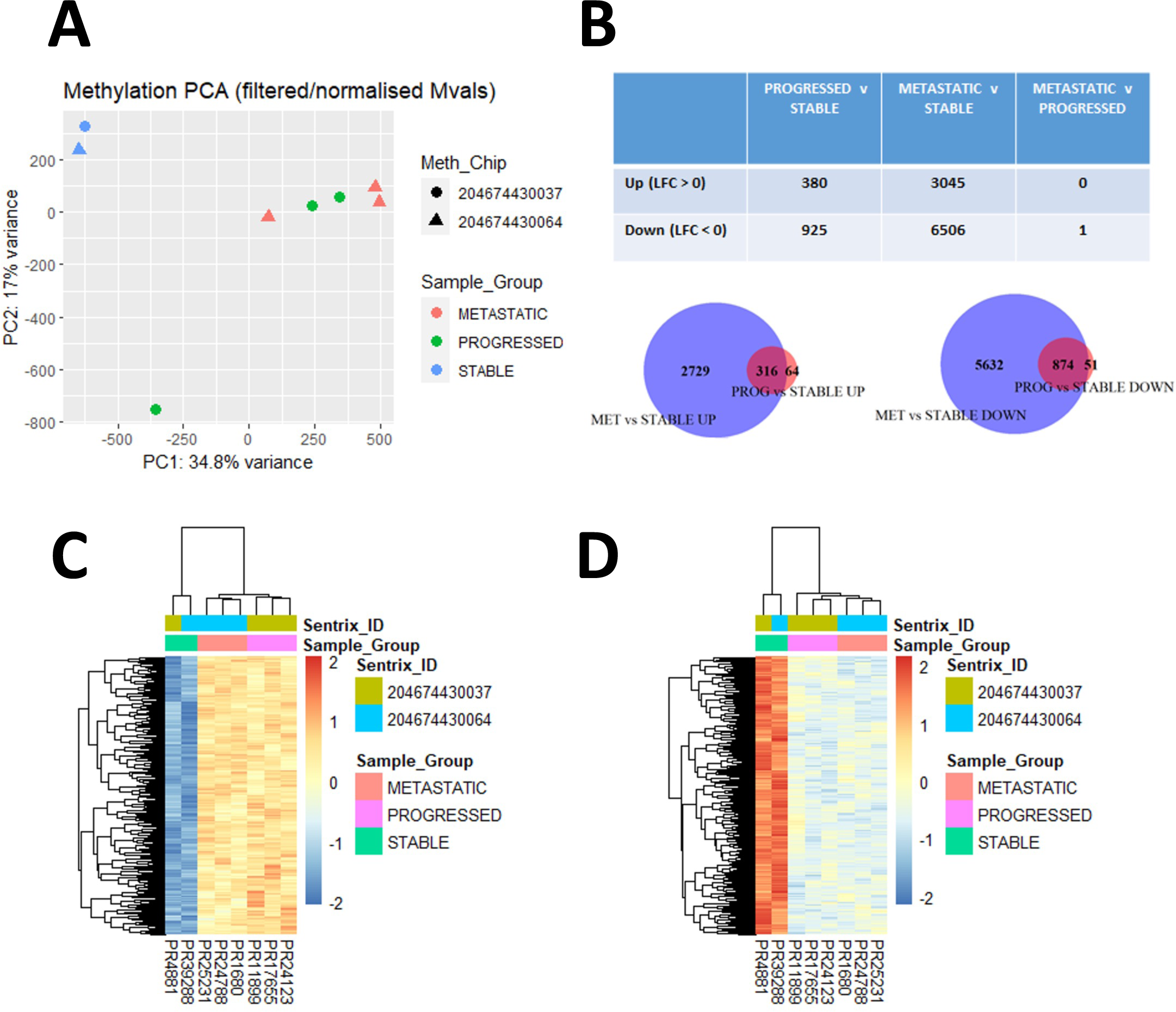
DNA methylation analysis of the cohort. Principal Component Analysis (PCA, post-normalization and filtering) demonstrated some separation of stable cases from the progressed and *de novo* metastatic cases by Principal Component 1. (**A**). Analysis of differential DNA methylation (post-normalization and filtering) demonstrated minimal differential methylation (adjusted p-value < 0.05) between progressed and *de novo* metastatic cases, and directionally coherent overlap between progressed versus stable cases, and between *de novo* metastatic versus stable cases, bases on the differentially methylated probes (Mvals) (**B**). A heatmap demonstrated overlap of differentially methylated probes from **B**, increased (LFC > 1) (**C**) and decreased (LFC < -1) (**D**) in progressed and *de novo* metastatic versus stable cases.

Analysis of differentially methylated regions (DMRs) was performed to identify whether the differentially methylated probes correspond to known genes. Over 600 DMRs were identified in a comparison of sPCa versus pPCa or mPCa cases, including genes of biological relevance in PCa such as *GNAS* and *AR* (Figures S14-S15). The DMR plots demonstrate pPCa and mPCa cases had similar mean methylation patterns, distinct from sPCa cases. The coherently differentially methylated probes (**Figure 5B**) were mapped to their genomic location to identify DMRs, to explore whether they were specific to individual chromosomes. The outer track of **Figure 6A** demonstrates the chromosomal location of hypomethylated (blue) and hypermethylated (red) regions, with the largest peak occurring at chromosome 19. The Rainfall plot middle track of **Figure 6A** shows the genomic coordinates of each region, with the y-axis corresponding to the minimum distance to neighbouring regions, demonstrating clustering of differentially methylated probes at Chromosome 19. To investigate whether the corresponding genes are part of specific pathways, an over-representation analysis was performed, which identified pathways associated (*padj*<1x10^-14^) with RNA pol II cis-regulatory region sequence-specific DNA binding, DNA binding transcription factor activity (GO) and Herpes Simplex 1 infection (KEGG) (Figures S16-S19). To explore the relationship between differential methylation and gene expression, the coherently hypomethylated probes were filtered between pPCa and mPCa cases, versus sPCa cases (**Figure 5B**), to identify potential association with gene promoters. This data was integrated with 3’RNAseq (rlog normalized) expression data for the corresponding 84 genes to generate combined methylation and expression heatmaps and density plots. Unsupervised hierarchical clustering of the differentially methylated probes corresponding to promoter regions demonstrated clustering of sPCa cases versus pPCa and mPCa cases (**Figure 6B, Top Panel**). Most of the probes corresponded to open chromatin regions and DNase I hypersensitivity sites. The methylation density heatmap (**Figure 6B, Second Panel**) visualized the distribution of methylation (mVals), and this was similar and compact for pPCa and mPCa cases. 3’RNAseq expression of genes corresponding to the promoter-related probes (**Figure 6B, Third Panel**) demonstrated variation in the corresponding gene expression, particularly in the sPCa samples. Density plots of gene expression demonstrated that the expression of genes corresponding to hypomethylated promoters was more tightly distributed in the pPCa and mPCa samples, whereas gene expression in the sPCa samples (where the promoter-associated probes were comparatively hypermethylated) showed greater variation (**Figure 6B, Bottom Panel**). These findings were similar in the RNA expression of differentially methylated genes in the full cohort (Figure S20). Over-representation analysis of hypo- and hypermethylated genes, performed using two separate platforms (enrichr and gprofiler), to compare sPCa cases versus pPCa and mPCa cases, identified pathways associated with histone H3 lysine 4 trimethylation (H3K4me3), histone H3 lysine 27 trimethylation (H3K27me3), and Polycomb Repressive Complex 2 (PRC) (Figure S21). To explore the relationship between EZH2 expression and expression of EZH2-regulated genes in sPCa versus pPCa and mPCa cases, it was observed that a set of genes previously identified as being regulated by EZH2^27^ were DEGs between these clinical groups of cases (*padj*<0.05) (Figure S22).

**Figure 6.**
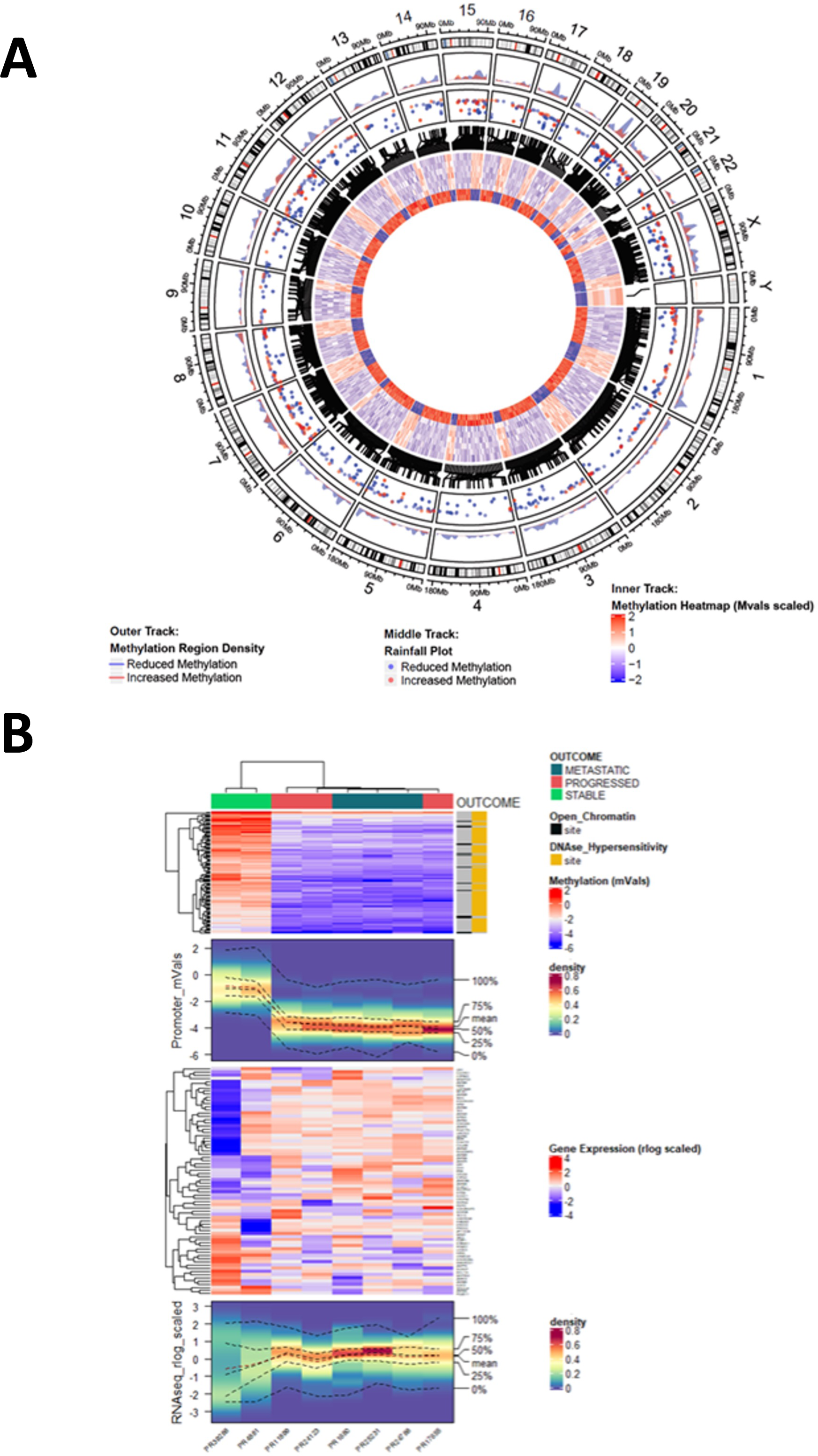
Analysis of coherently differentially methylated probes versus their genomic location and versus gene expression data. A Differential Methylation Density plot of coherently differentially methylated regions between progressed and *de novo* metastatic cases versus stable cases. (**A**). The outer track density plot demonstrated the fraction of the genomic window covered by differentially methylated regions in progressed and *de novo* metastatic cases versus stable cases (hypo-methylated regions in blue, hyper-methylated regions in red). The middle track rainfall plot demonstrated the genomic coordinates of each region (the y-axis corresponding to the minimum distance to the neighboring region). The inner track circular heatmap demonstrated coherently differentially methylated probes (Mvals scaled). The outer 3 lanes correspond to *de novo* metastatic cases; the middle 3 lanes correspond to progressed cases; the inner 2 lanes correspond to stable cases. A combined promoter methylation and gene expression analysis is shown in (**B**). The top panel methylation heatmap demonstrated promoter-associated probes hypomethylated in progressed and *de novo* metastatic cases versus stable cases. Open chromatin regions (OCRs) and Dnase I hypersensitivity sites (DHSs) are annotated, demonstrating promoter-associated CpG sites as either DHSs or OCRs. The second panel methylation density heatmap visualizes the distribution of Mvals, with clustering as per the top panel. The third panel demonstrated the expression (Quantseq) of genes corresponding to CpGs in the upper two panels; gene expression is normalized (rlog – DESeq2) and scaled. The fourth and final panel is a gene expression density heatmap displayed as density of Quantseq expression values.

Whilst this proof-of-concept study did not specially aim to identify a gene signature prognostic for clinical outcome post-RT, a set of genes was demonstrated to be prognostic for post-RT outcome in a large external dataset (Figure S23). This set of genes was distinct from gene sets previously described in the literature and warrants further investigation.

## Discussion

This proof-of-concept study tested the possibility that orthogonal genomic analyses (3’RNAseq, nanoString and DNA methylation) can identify molecular features associated with PCa progression in the baseline historical pre-treatment FFPE biopsy samples with long-term clinical follow-up. This unlocks the potential to investigate powerful large clinical cohorts with long-term follow-up, such as the ProtecT cohort^1^. It raises the potential to utilize molecular features of prostate biopsy features in clinical decision making, to aid risk-stratification and personalized medicine approaches in the post-diagnostic space. Both retrospective studies of large historical cohorts, and prospective studies to investigate the added value of this approach in the clinical setting, are now warranted.

Transcriptional signatures prognostic of metastatic recurrence have been described in PCa, and some of these have been validated in patients treated with RT^5-11^. However, most of these studies, except for the ‘hypoxia signature’, describe signatures derived from radical prostatectomy specimens rather than from pre-treatment prostate biopsies from patients undergoing RT. DNA methylation is altered in PCa development and progression, however whilst prognostic biomarkers have been developed this approach has not been specifically used for RT-treated patients^16,17,28^.

Large clinical datasets, such as the DECIPHER GRID project (NCT02609269) which contains transcriptomic information from 20,000 patients, are required to reliably identify molecular features associated with disease behavior. The results of this proof-of-concept study highlight the opportunity for future larger studies to obtain molecular data from historical diagnostic prostate biopsy samples as a method of identifying key molecular features of high-risk disease. It is demonstrated herein that 3’RNAseq, nanoString, and methylation analysis each have the capacity to achieve this, and to identify baseline molecular features associated with development of metastatic PCa.

One sample in this analysis failed 3’RNAseq, with all other analyses achieving useable data. One sample achieved fewer counts than others (500,000 versus ∼8 million), however this sample was included in the analysis for several reasons. First, 3’RNAseq only measures 3’ transcripts and produces counts for a whole transcript, whereas standard RNA sequencing measures multiple fragments across the same gene transcript. Second, differential expression analysis tools can account for differing library sizes. Third, the analysis was also performed with this sample excluded, with minimal change to the results. In addition, to explore the impact of mapping techniques, the analysis was repeated utilizing Subread Aligner versus STAR with this sample both included and excluded, achieving similar results.

The 3’RNAseq DEGs analysis demonstrated several broad similarities in gene expression between pPCa cases and mPCa cases, both in terms of the PCA, the pattern, number and directional coherence of DEGs between groups, and the non-hierarchical clustering analysis. Two groups of samples (pPCa and mPCa) were clearly distinct from sPCa cases. This observation suggests that there is shared underlying molecular biology between pPCa and mPCa cases, that can be identified in archival FFPE pre-treatment prostate biopsies using these techniques. This may potentially be explained by performance limitations of radiological staging at the time of original diagnosis of the historical cases with long-term follow-up selected for this proof-of-concept study. For example, radionuclide bone scans were used for baseline staging as part of risk-stratification, rather than the more recently developed MRI marrow or PSMA PET CT scans. It is possible that the original TNM staging of these cases, using imaging modalities available at the time, led to under-staging of some cases. The cases with pPCa may have had undetectable micro-metastases at the time of original diagnosis and treatment, thus accounting for these baseline prostate biopsy samples having similar molecular features compared to the baseline mPCa cases, with both of these two groups of patient samples being clearly distinct from sPCa cases. Nevertheless, it is interesting that such differences can be identified in the relatively small amount of genomic material available from archival FFPE prostate biopsy samples. This raises the exciting possibility that the added value of molecular analysis approach could be taken forward for evaluation in prospective clinical studies investigating the utility of this approach in treatment decision making. This would aim to improve risk stratification and clinical outcomes for patients, and warrants investigation in future studies.

The similarities in RNA expression in pPCa and mPCa cases, compared with sPCa cases, observed in the 3’RNAseq analysis, were also generally observed in the nanoString analysis using PCA and gene expression tools. However, in the nanoString analysis there is incomplete separation of the sPCa cases, with one mPCa case clustering with the sPCa cases. This may potentially be explained by the nanoString platform being a targeted panel containing significantly fewer genes than those used for 3’RNAseq (∼800 versus >20,000).

Moreover, the two techniques have different mechanisms of action, with nanoString utilizing reporter probes to hybridize mRNA, and 3’RNAseq sequencing the 3’ end of transcripts, which could be potentially affected by alternative splicing. The Spearman correlation between 3’RNAseq (rlog normalized) expression and nanoString (log normalized) expression for the subset of nanoString genes was strong at 0.68, and per-sample correlations utilising different normalization techniques for 3’RNAseq were similar.

Whilst the demonstrated DEGs (*p*<0.05) in the 3’RNASeq analysis did not achieve statistical significance on correction for multiple testing using the NSolver^TM^ nanoString analysis, the results presented herein are scaled values of normalized expression, which ought to be considered to broadly support the findings of the 3’RNASeq analysis rather than be utilized in isolation. The use of these orthogonal RNA sequencing techniques demonstrates similar findings across these transcriptional analysis platforms comparing pPCa (and mPCa) cases versus sPCa cases in this proof-of-concept study.

Over-representation analysis identified Gene Ontology pathways associated with spindle pole and centrosome function to be increased in pPCa and mPCa cases, versus sPCa cases, and centrosome pathways were also identified on ssGSEA, potentially due to increased mitotic activity in those samples. An RNA expression-based CCP score is independently prognostic of metastatic progression after RT, although there is minimal overlap of genes between genesets^7, 18^. Increased expression of spindle pole genes in pPCa cases could be due to increased reliance on the spindle assembly checkpoint, due to loss of other cell checkpoints^29-30^. Increased expression of centrosome pathway genes could be due to centrosome amplification in primary tumor samples of patients who develop metastatic disease. Previous studies have demonstrated *in situ* centrosome loss in primary PCa samples^31^, however in PCa cell lines centrosome amplification has been observed in PCa cell lines with increased metastatic behavior^32^. Both observations may potentially be related to increased chromosomal instability and aneuploidy.

Four genes (*GNAS*, *ETV1*, *COL2A1* and *HDAC5*) were observed to be coherently and significantly differentially expressed between sPCa and pPCa cases using both 3’RNAseq (*padj*<0.05) and nanoString platforms (less stringent *p*<0.2), however only *COL2A1* and *ETV1* were prognostic upon application of a cox proportional hazards model. An integrative clinical genomic study previously demonstrated *GNAS* to be one of the most frequently mutated genes in advanced PCa^31^. *ETV1* is a transcription factor frequently overexpressed in aggressive PCa via a chromosomal translocation with androgen-responsive promoters^33^, and *ETV1* has been demonstrated to initiate PCa tumorigenesis in concert with the JMJD2 histone demethylase^34^. *COL2A1* is a candidate PCa risk gene^35^, and the *HDAC5* histone deacetylase gene is frequently downregulated or deleted in PCa, resulting in increased H3K27 acetylation and impaired RB-mediated repression of cell cycle-related pro-oncogenic genes^36^. In this proof-of-principle study dataset, *HDAC5* expression is reduced in sPCa versus pPCa cases analyzed with both RNA expression techniques. Analysis of an external dataset of radiotherapy-treated patients^11^ demonstrates that these four genes (*GNAS*, *ETV1*, *COL2A1* and *HDAC5*) are prognostic for biochemical progression-free survival, though not for metastatic progression-free survival.

Features obtained from ten previously described metastatic signatures, from various other cancer types, were observed be differentially expressed upon ssGSEA analysis between sPCa cases and pPCa and mPCa cases. This clear separation of samples by PCA using genes specifically associated with metastasis identifies similarities in samples from patients with pPCa and mPCa, distinct from sPCa cases. This was further demonstrated utilizing the top 50 DEGs (of which 36 are represented) from a comparison of sPCa versus pPCa cases in a large external dataset^11^. It is noteworthy that there is no overlap between these genes and those in the Prolaris, Oncotype, Decipher, Metastatic Assay and Prostate Hypoxia genelists. Separation of the cohort by the median expression into HIGH and LOW groups was observed to be prognostic for both biochemical and metastatic progression-free survival, whilst these genes were more prognostic than would be expected from a random set of genes and selected cancer signatures. Four genes (*CDC25A*, *OLR1*, *CDON* and *DDX39B*) were demonstrated to be independently prognostic, and each of these has been previously reported to be associated with PCa progression and/or metastasis^37-40^. Whilst it is a limitation of this proof-of-concept study that the analysis is underpowered for biomarker discovery, these results demonstrate that 3’RNAseq analysis of historical FFPE prostate biopsy samples with long-term follow-up can identify biologically relevant genes which can be validated using a large external dataset.

DNA hypomethylation has been described in PCa, and recent studies in advanced disease have identified specific areas of increased hypomethylation during progression from benign tissue to localised PCa to metastatic disease^41^. The findings in this study are consistent with this phenomenon, with similar patterns of methylation being observed in pPCa and mPCa cases, and increased methylation in mPCa versus sPCa cases, compared to pPCa versus sPCa cases.

Methylation analysis demonstrated separation of sPCa cases and pPCa/mPCa cases using both PCA and DEGs analysis, with the majority of differential gene methylation being coherent and in the same direction, accepting that a limitation of this proof-of-concept study is the relatively small number of patient samples. Previous studies have demonstrated similar methylation patterns for primary and metastatic tumor samples from the same patient^42^, supporting our observation that methylation characteristics of metastatic disease can be found in primary tumor biopsies. These DNA methylation results provide additional evidence beyond the 3’RNAseq and nanoString data of biological similarities in baseline samples from patients with pPCa and mPCa at both the methylomic and transcriptomic level.

The methylation technique used in this study, using 850K methylation probes, has the granularity to identify multiple areas of methylation within a single gene. We identified specific differentially methylated DNA regions which map to genes of known PCa biological relevance, including *AR* and *GNAS*. *AR* hypomethylation has been observed in metastatic PCa^42-43^, and *GNAS* was hypomethylated and upregulated in pPCa/mPCa samples versus sPCa samples. The relationship between DNA methylation and gene expression is complex, with increased gene expression being associated with hypermethylation^44^. Our analysis of differential methylation plots demonstrated that pPCa and mPCa samples were hypomethylated at the start of genes, and hypermethylated in other regions, compared with sPCa cases. In the case of *GNAS*, we also observed areas of differential methylation within the same gene^44^.

The observation that the greatest focus of hypomethylation in pPCa and mPCa samples versus sPCa samples was on chromosome 19 is interesting, given that RNA Pol II–associated chromatin interactions have been identified as determinants of transcriptional regulation in PCa^45^. Moreover, there is significant over-representation, on pathway enrichment analysis, of associated genes at the site of peak differential methylation density on chromosome 19. It is also noteworthy that RNA Pol II interactions frequently involve H3K4m3 and/or H3K27 acetylation marks^45^. The over-representation analysis of differentially methylated probes identified pathways associated with H3K4m3, H3K27m3, and PRC2, and these pathways appeared in both hyper- and hypomethylated genes, demonstrating the complexity of methylation events in different areas of the same gene. The Polycomb Group protein Enhancer of Zeste 2 (EZH2) in the PRC2 complex catalyses H3K27m3 on target gene promoters, and EZH2 function has previously been associated with metastatic PCa progression, and with metastatic progression post-RT^25,46-49^. We observed several genes associated with EZH2 to be significantly differentially expressed between pPCa/mPCa cases and sPCa cases.

## Conclusion

This study demonstrates the potential for molecular analysis of diagnostic baseline prostate biopsy samples as a tool to characterize PCa beyond the current method of risk-classification, ahead of potential curative or systemic therapy, with added value in terms of identifying patients with occult micro-metastatic disease. This warrants further investigation in both retrospective studies using larger cohorts, and prospective studies designed to investigate the use of these molecular techniques in the clinic. Taken together, the results of this proof-of-concept study demonstrate that we can now unlock the potential wealth of information that can be gained from molecular analyses of powerful large historical cohorts with baseline FFPE prostate biopsy samples and associated long-term clinical follow up. This approach may aid future risk-stratification and treatment selection in the post-diagnostic space for men with this common malignancy^50^.

## Supporting information

Supplementary Package Summary

---

**Table S1.** Baseline clinicopathological characteristics of the cohort.

| Group | T | N | M | Primary<br>Gleason | Secondary<br>Gleason | Gleason<br>Sum | Grade<br>Group | PSA<br>(ng/ml) | Follow-up<br>(years) | Risk Group/<br>Metastatic |
| --- | --- | --- | --- | --- | --- | --- | --- | --- | --- | --- |
| Stable | T1c | N0 | M0 | 3 | 3 | 6 | 1 | 7.2 | 8.17 | Low |
| Stable | T2 | N0 | M0 | 4 | 5 | 9 | 5 | 8.7 | 6.75 | High |
| Stable | T2c | N0 | M0 | 3 | 4 | 7 | 2 | 1.2 | 6.75 | High |
| Stable | T3b | N0 | M0 | 4 | 3 | 7 | 3 | 11 | 8.17 | High |
| Stable | T3b | N0 | M0 | 3 | 4 | 7 | 2 | 80.3 | 8.25 | High |
| Progressed | T2c | N0 | M0 | 4 | 3 | 7 | 3 | 52 | 6.58 | High |
| Progressed | T2a | N0 | M0 | 5 | 4 | 9 | 5 | 4.7 | 3.75 | High |
| Progressed | T2c | N0 | M0 | 4 | 3 | 7 | 3 | 43.7 | 6.75 | High |
| Progressed | T1c | N0 | M0 | 4 | 3 | 7 | 3 | 2.5 | 4.33 | Salvage RT |
| Progressed | T3b | N0 | M0 | 4 | 3 | 7 | 3 | 6.4 | 3.75 | High |
| Metastatic | T2c | N0 | M1c | 3 | 4 | 7 | 2 | 18.7 | 5.66 | Metastatic |
| Metastatic | T4 | N0 | M1 | 4 | 5 | 9 | 5 | 52.4 | 5.08 | Metastatic |
| Metastatic | T2c | N0 | M1b | 4 | 5 | 9 | 5 | 149.5 | 4.17 | Metastatic |
| Metastatic | T3 | N0 | M1b | 4 | 5 | 9 | 5 | 19.1 | 2.42 | Metastatic |
| Metastatic | T4 | N1 | M1b | 4 | 4 | 8 | 4 | 15856.6 | 2.33 | Metastatic |
| Metastatic | T3b | N0 | M1b | 4 | 5 | 9 | 5 | 1524.6 | 1.42 | Metastatic |
| Metastatic | T3b | N0 | M1b | 4 | 5 | 9 | 5 | 31.4 | 1.58 | Metastatic |
| Metastatic | T3a | N1 | M1b | 4 | 4 | 8 | 4 | 513.4 | 1.0 | Metastatic |

**Figure S1.**
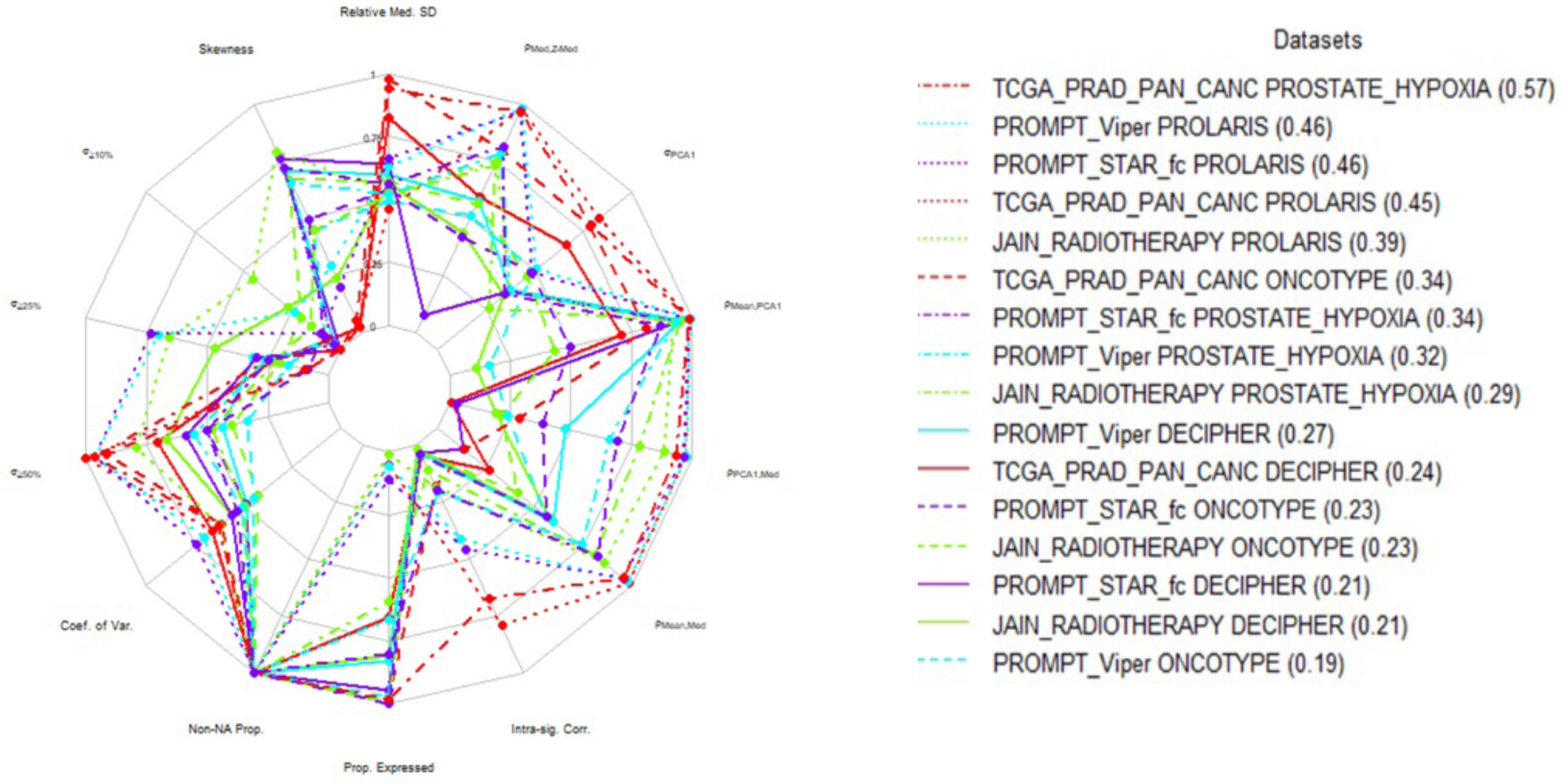
Signature Summary scores produced by SigQC for previously validated signatures in prostate cancer (Prostate Hypoxia, Prolaris, Oncotype and Decipher) in the PROMPT dataset compared to TCGA Prostate PanCancer and Jain Radiotherapy datasets. Overall signature performance is comparable in the PROMPT dataset compared to previously published datasets.

**Figure S2.**
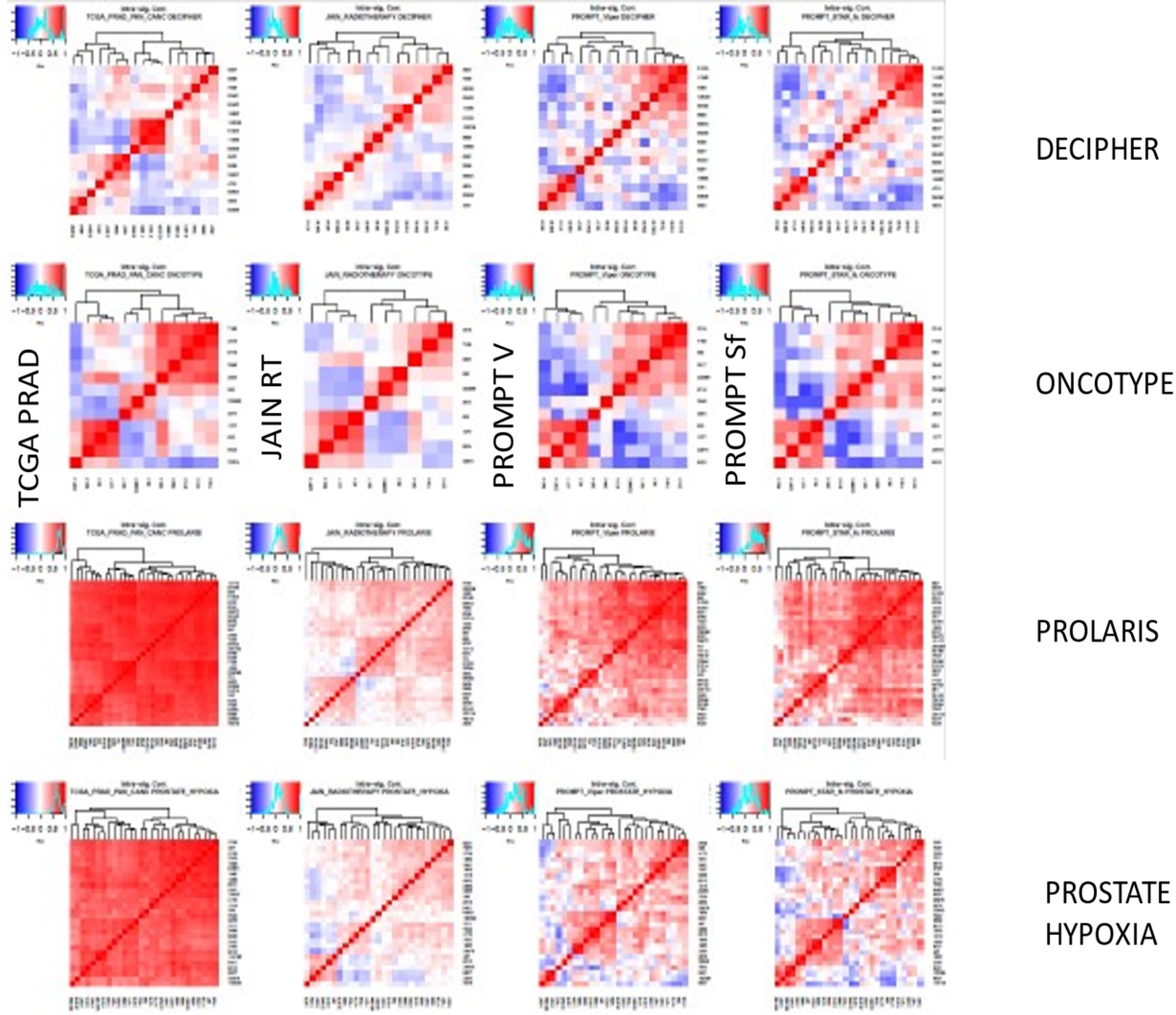
Intra-signature correlation generated by SigQC for previously validated signatures in prostate cancer (Prostate Hypoxia, Prolaris, Oncotype and Decipher) in the PROMPT dataset compared to TCGA Prostate PanCancer and Jain radiotherapy datasets. Intra-Signature correlation is comparable in the PROMPT dataset compared to previously published datasets.

**Figure S3.**
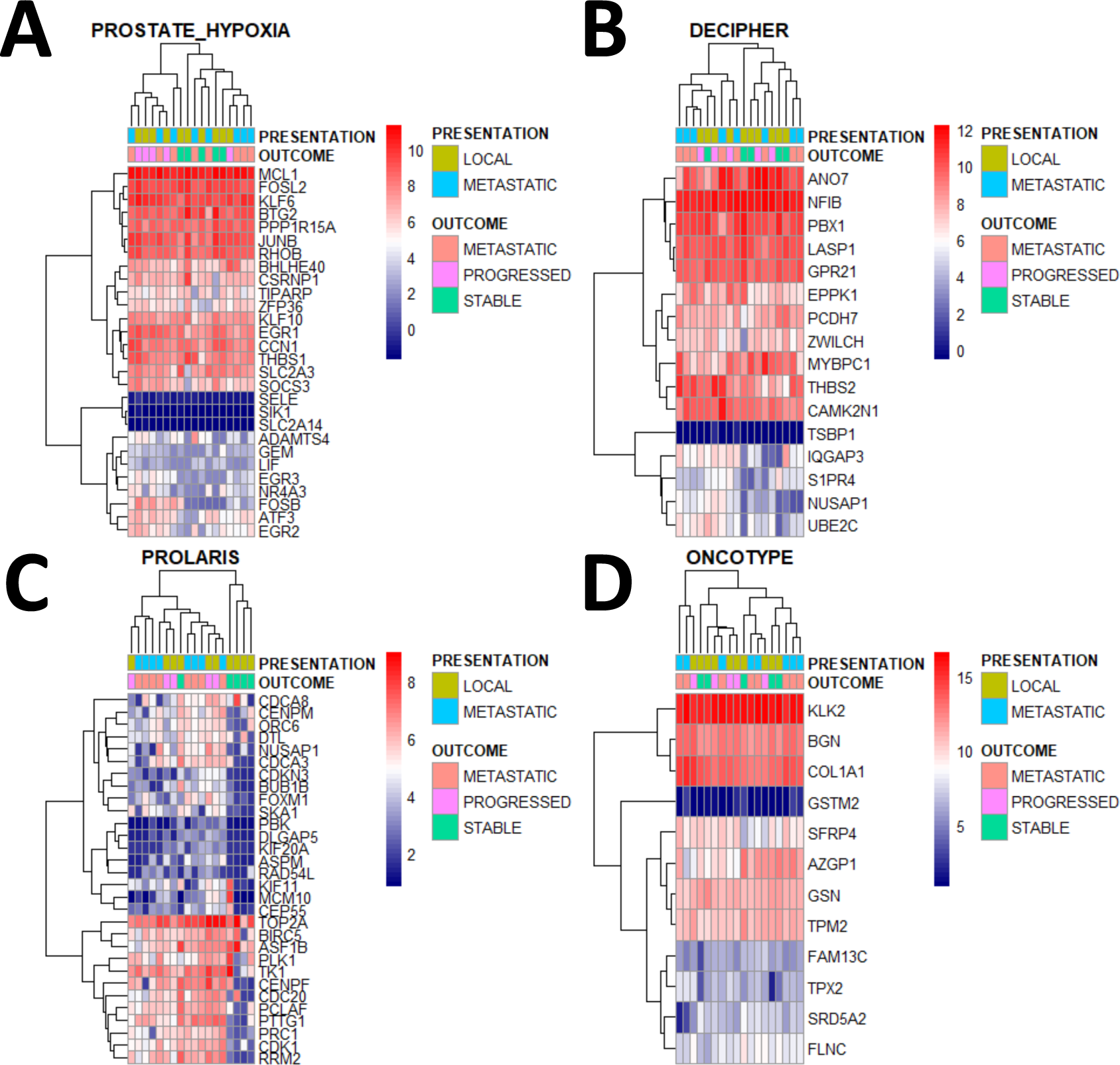
Signature expression heatmaps. Expression of previously validated prostate cancer signatures in PROMPT dataset; Prostate Hypoxia (**A**), Decipher (**B**), Prolaris (**C**), Oncotype (**D**).

**Figure S4.**
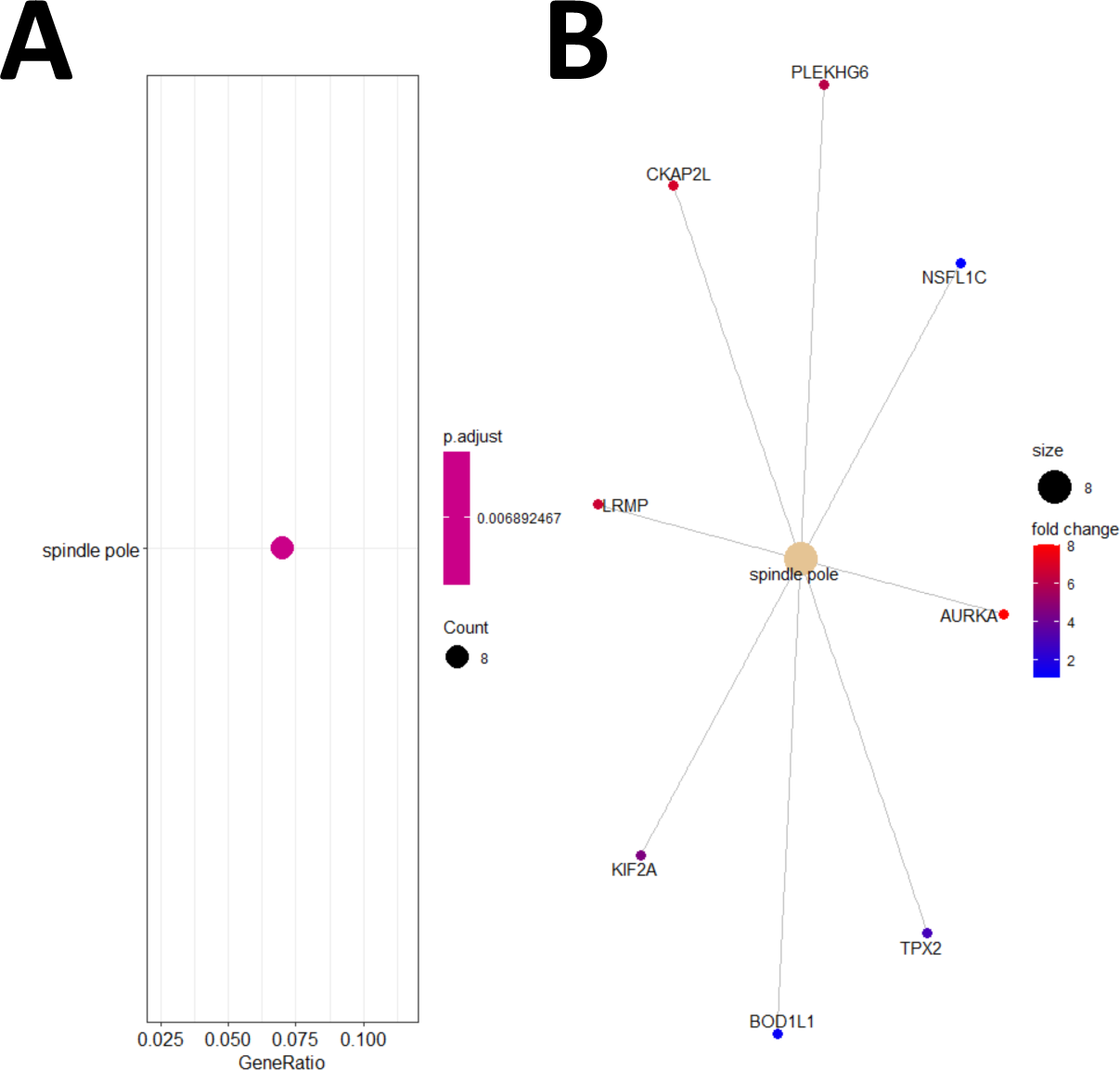
Over-representation analysis of genes increased in progressed versus stable prostate cancer patients post-RRT (padj <0.01) (**A**). A network plot visualizing the enriched genes (**B**).

**Figure S5.**
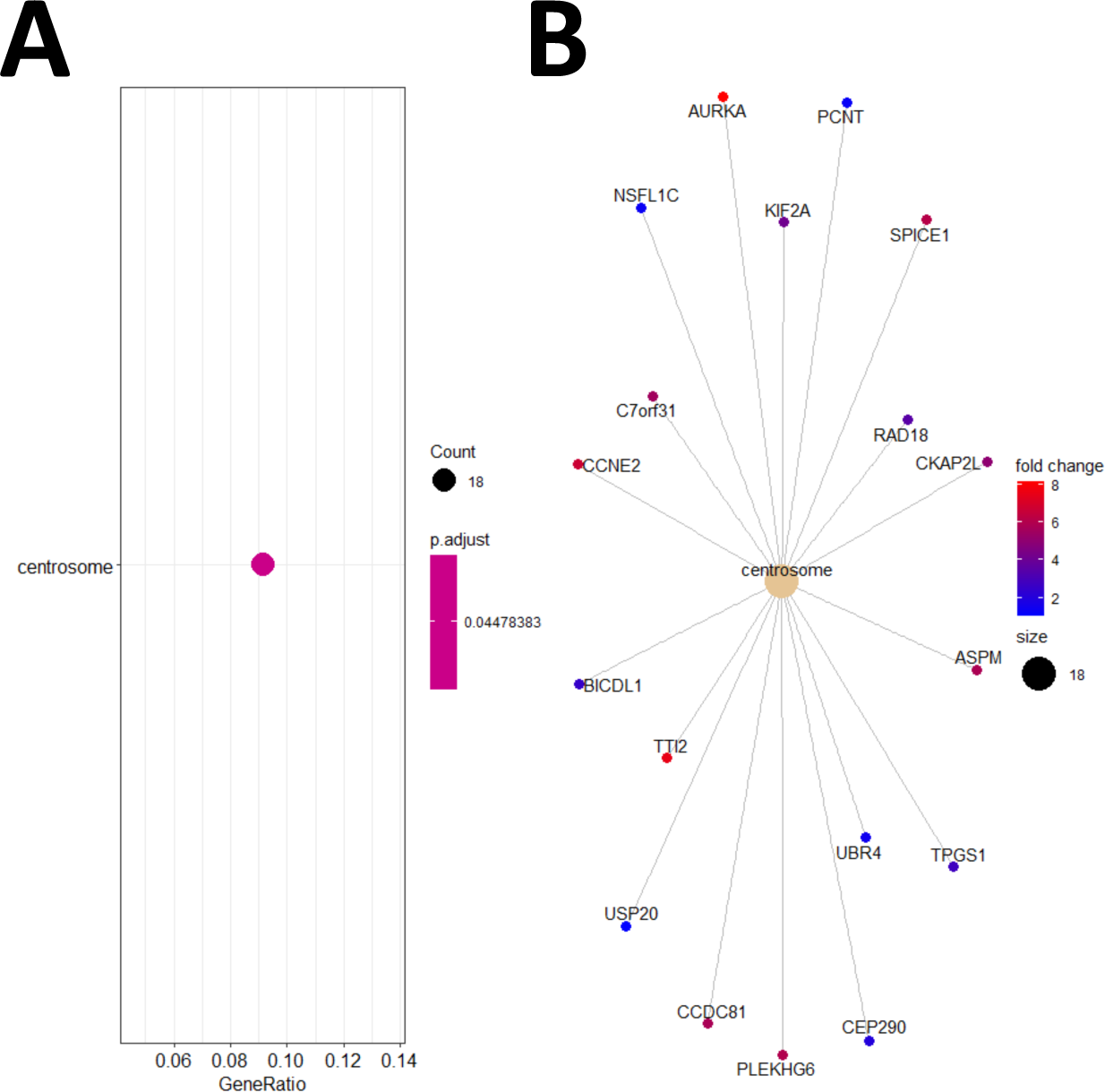
Over-representation analysis of genes increased in metastatic versus stable prostate cancer patients post-RRT (padj <0.05) (**A**). A network plot visualizing the enriched genes (**B**).

**Table S2.** Overlap of significantly differentially expressed genes between progressed and stable samples in both Quantseq (padj <0.2) and Nanostring (p<0.05) analysis. Genes increased in progressed versus stable cases in Quantseq and Nanostring include GNAS, ETV1, COL2A1. HDAC5 expression is decreased in progressed versus stable cases in Quantseq and Nanostring. LogFC - Log Fold Change; lfcSE - standard error; padj - adjusted p-value (**A**). A per gene univariable cox proportional hazards analysis was performed for these genes using the Jain Radiotherapy dataset (**B**).

|  |  |  |  |  |  |  |  |  |  |  |  |  |
| --- | --- | --- | --- | --- | --- | --- | --- | --- | --- | --- | --- | --- |
| Quantseq |  |  |  |  |  |  |  |  |  |  |  |  |
|  | baseMean | logFC | lfcSE | stat | pvalue | padj | ENTREZ | SYMBOL |  |  |  |  |
| GNAS | 3424.709 | 0.657159 | 0.204451 | 3.214258 | 0.001308 | 0.039557 | 2778 | GNAS |  |  |  |  |
| HDAC5 | 566.0214 | -1.28296 | 0.332002 | -3.86431 | 0.000111 | 0.006872 | 10014 | HDAC5 |  |  |  |  |
| ETV1 | 614.1381 | 2.874883 | 1.113311 | 2.582281 | 0.009815 | 0.142602 | 2115 | ETV1 |  |  |  |  |
| COL2A1 | 505.9484 | 7.520908 | 1.50439 | 4.999306 | 5.75E-07 | 0.000125 | 1280 | COL2A1 |  |  |  |  |
| NANOSTRING |  |  |  |  |  |  |  |  |  |  |  |  |
| hgnc_sym | logFC | std.error.. | Lower.cor | Upper.cor | Linear.fold | Lower.cor | Upper.cor | P.value | BY.p.value | method | Gene.sets | probe.ID |
| GNAS | 1.12 | 0.157 | 0.808 | 1.42 | 2.17 | 1.75 | 2.68 | 5.60E-05 | 0.0937 | lm.nb | Driver Gen | NM_080425.1:1910 |
| HDAC5 | -0.393 | 0.0998 | -0.588 | -0.197 | 0.762 | 0.665 | 0.872 | 0.00432 | 0.544 | Wald | Chromatin | NM_005474.4:3160 |
| ETV1 | 3.6 | 1.1 | 1.45 | 5.75 | 12.1 | 2.74 | 53.8 | 0.00946 | 0.962 | lm.nb | Transcript | NM_004956.4:1719 |
| COL2A1 | 2.24 | 1.38 | -0.461 | 4.95 | 4.73 | 0.726 | 30.8 | 0.143 | 1 | Wald | PI3K | NM_001844.4:4745 |

B
| Variable | Beta | StandardError | Z | P | LRT | Wald | LogRank | HR | HRlower | HRupper | SYMBOL |
| --- | --- | --- | --- | --- | --- | --- | --- | --- | --- | --- | --- |
| ENSG00000139219 | 0.190489 | 0.08881475 | 2.144795 | 0.031969 | 0.048235 | 0.031969 | 0.030404 | 1.209842 | 1.01655118 | 1.43988482 | COL2A1 |
| ENSG00000108840 | 0.298047 | 0.500965996 | 0.594945 | 0.55188 | 0.555108 | 0.55188 | 0.551934 | 1.347225 | 0.50468132 | 3.59636105 | HDAC5 |
| ENSG00000087460 | 0.118691 | 0.319839477 | 0.371096 | 0.710566 | 0.709384 | 0.710566 | 0.710664 | 1.126022 | 0.60158793 | 2.10763219 | GNAS |
| ENSG00000006468 | 0.363658 | 0.076723086 | 4.739879 | 2.14E-06 | 6.31E-05 | 2.14E-06 | 2.26E-07 | 1.438582 | 1.23773566 | 1.67202021 | ETV1 |

**Figure S6.**
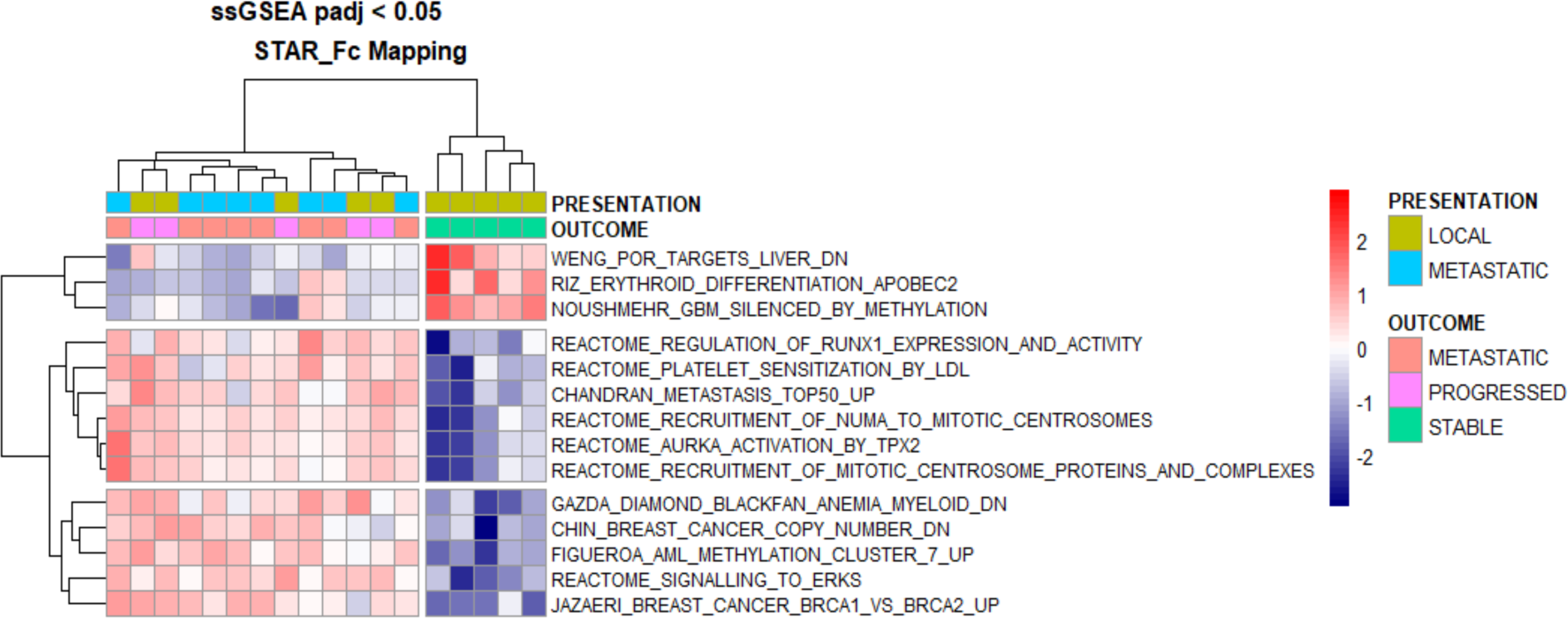
Single sample GeneSet Enrichment Analysis (ssGSEA) pathways analysis of progressed & metastatic cases versus stable cases post-RRT. ssGSEA identifies multiple pathways from MSigDB C2: curated gene sets, significantly different between progressed and metastatic cases versus stable cases post-RRT. 3 Pathways are enriched in stable cases, 11 pathways are enriched in progressed and metastatic cases.

**Figure S7.**
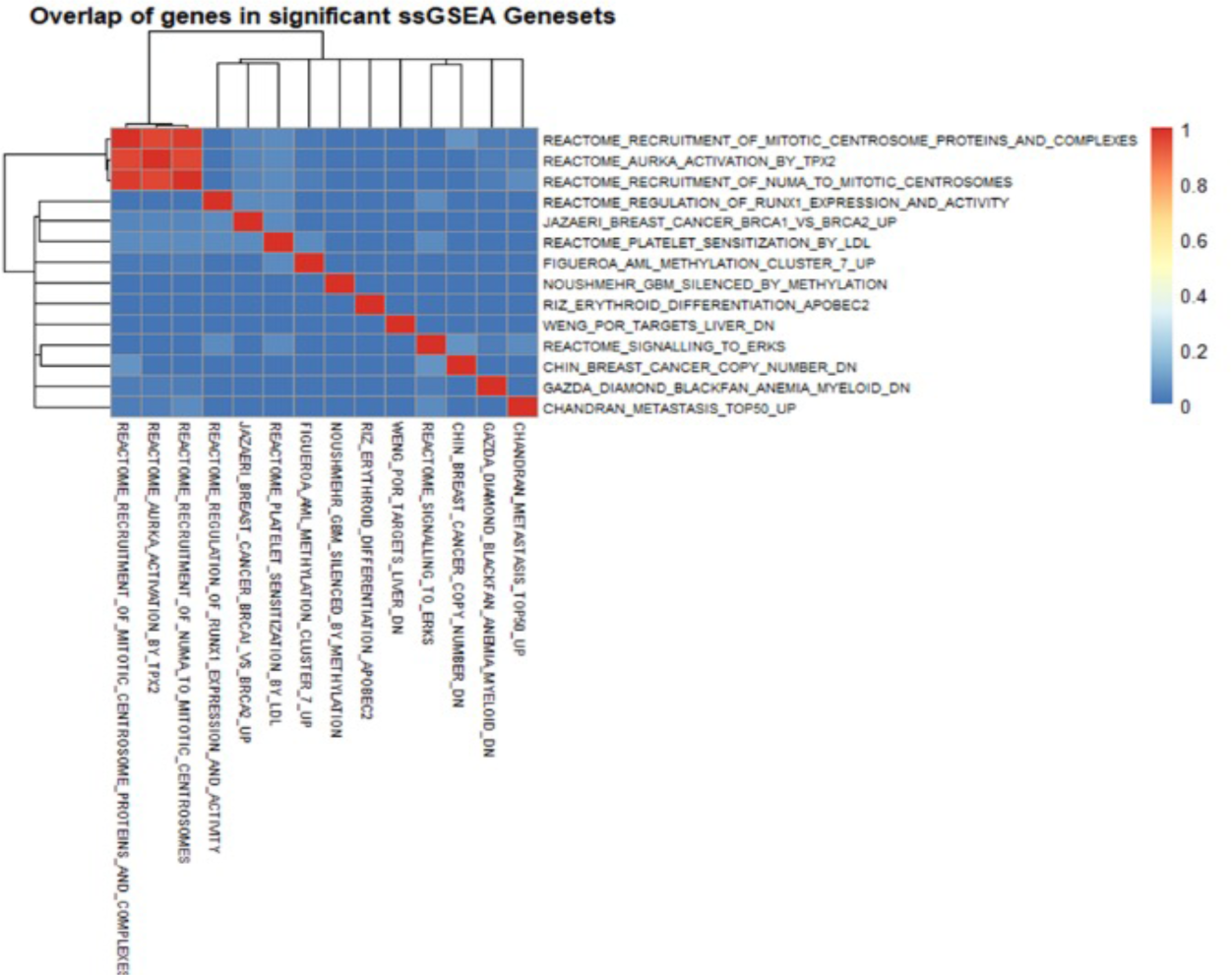
Overlap of genes in significant single sample GeneSet Enrichment Analysis (ssGSEA) pathways, comparing progressed and metastatic cases versus stable cases. Overlap is measured between 0 (0%) and 1 (100%).

**Table S3.** Guided single sample GeneSet Enrichment Analysis (ssGSEA) of MSigDB C2: curated metastasis gene set signatures in the Jain dataset. LogFC - Log Fold Change; AveExpr-Average Expression; t = t statistic; P.Value - p-value; adj.P.Val - adjusted p-value; B - B statistic.

|  | logFC | AveExpr | t | P.Value | adj.P.Val | B |
| --- | --- | --- | --- | --- | --- | --- |
| CHANDRAN_METASTASIS_TOP50_UP | 0.0432 | 0.940626 | 5.081351 | 5.59E-05 | 0.003184 | 1.864197 |
| BIDUS_METASTASIS_DN | 0.042476 | 0.519153 | 3.188963 | 0.004582 | 0.082714 | -2.3971 |
| CHANDRAN_METASTASIS_UP | 0.03397 | 0.634138 | 3.111228 | 0.00547 | 0.082714 | -2.5649 |
| TAVAZOIE_METASTASIS | -0.03039 | 0.024315 | -3.08515 | 0.005804 | 0.082714 | -2.62088 |
| TOMIDA_METASTASIS_UP | 0.033769 | 0.718665 | 2.921186 | 0.008403 | 0.095794 | -2.96901 |
| RAMASWAMY_METASTASIS_UP | 0.036273 | 0.644774 | 2.750608 | 0.012278 | 0.099163 | -3.32314 |
| LIAO_METASTASIS | 0.02087 | 0.497343 | 2.736128 | 0.012676 | 0.099163 | -3.35278 |
| PROVENZANI_METASTASIS_UP | 0.022527 | 0.572919 | 2.596877 | 0.017182 | 0.099163 | -3.63414 |
| LEE_METASTASIS_AND_RNA_PROCESSING_UP | 0.049333 | 0.680765 | 2.592328 | 0.017352 | 0.099163 | -3.64321 |
| PEDERSEN_METASTASIS_BY_ERBB2_ISOFORM_1 | -0.05147 | 0.104123 | -2.59114 | 0.017397 | 0.099163 | -3.64558 |

**Figure S8.**
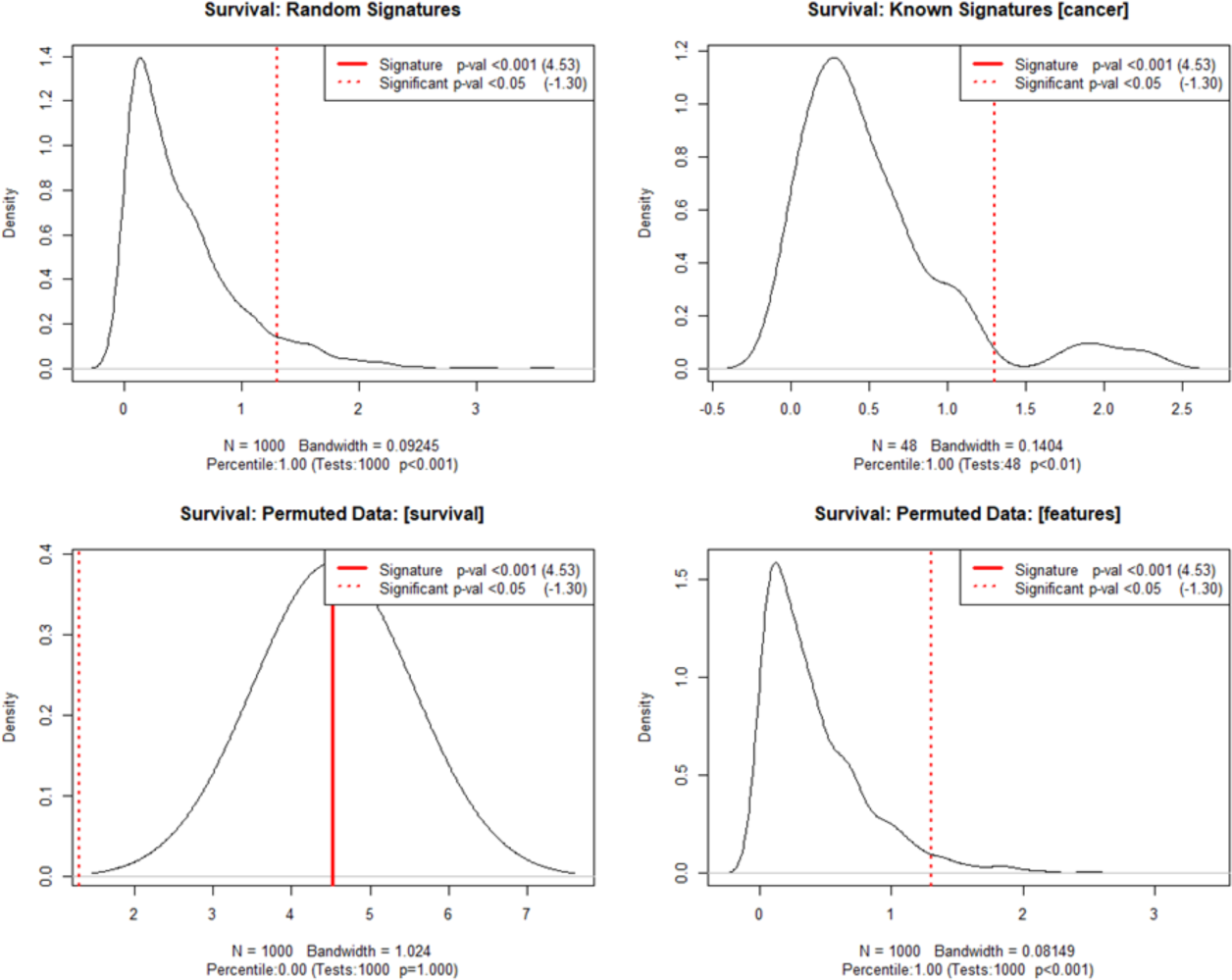
SigCheck comparison of random signatures, cancer signatures, and survival and feature permutations for biochemical progression-free survival in the Jain dataset. The vertical red dotted line shows where a “significant” result (p=0.05) would lie relative to the background distribution.

**Figure S9.**
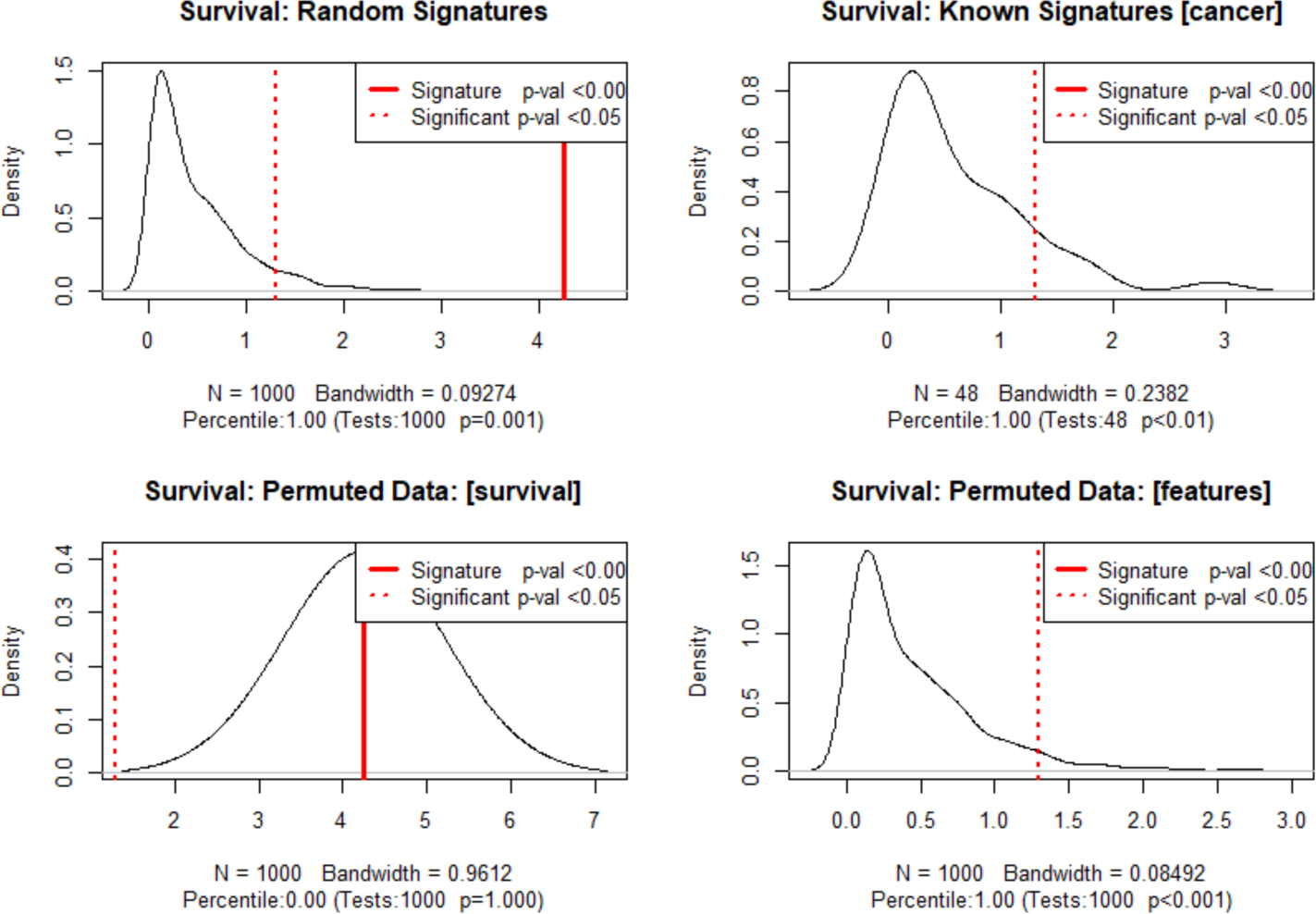
SigCheck comparison of random signatures, cancer signatures, and survival and feature permutations for metastatic progression-free survival in the Jain dataset. The vertical red dotted line shows where a “significant” result (p=0.05) would lie relative to the background distribution.

**Figure S10.**
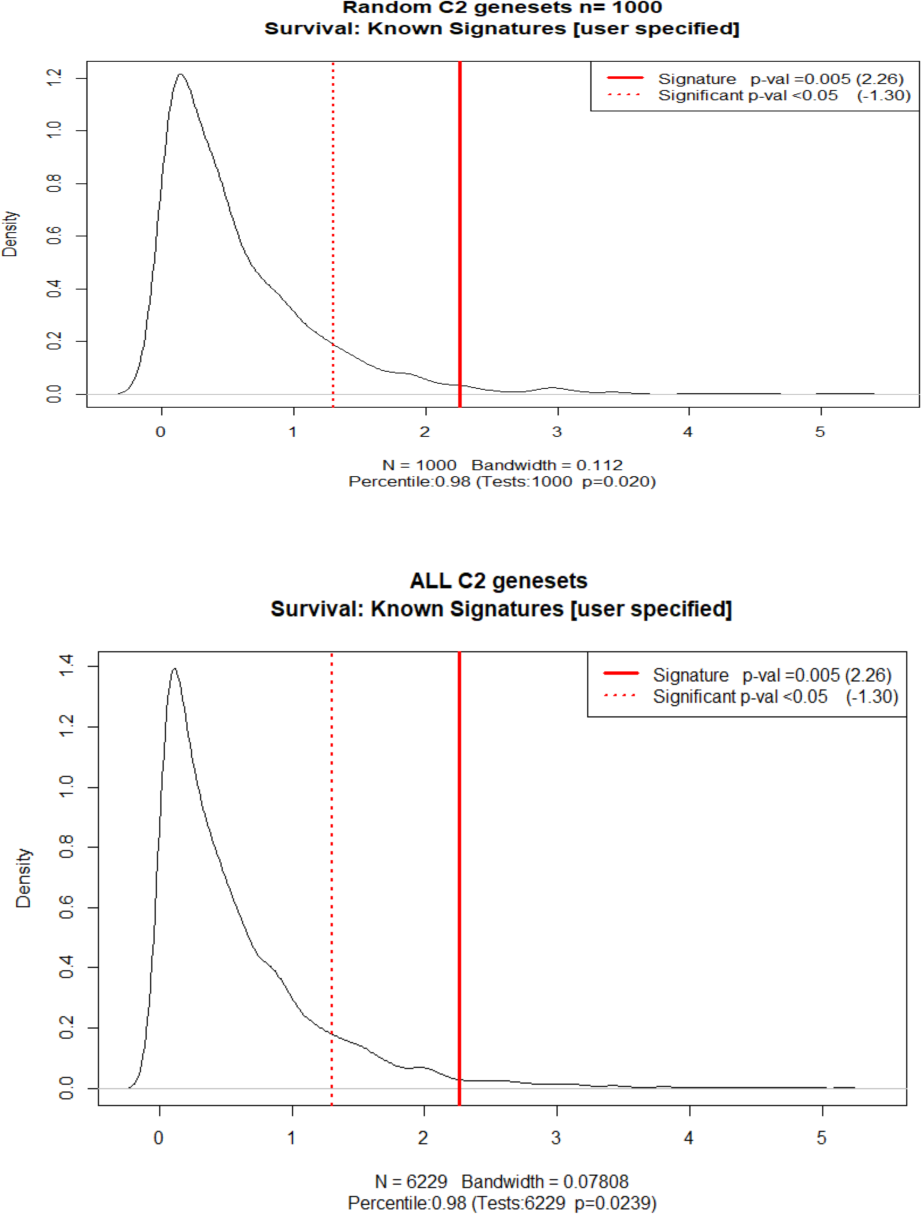
SigCheck comparison of subset and all curated cancer signatures (MsigDB) compared to PROMPT signature for metastatic progression-free survival in the Jain dataset. The vertical red dotted line shows where a “significant” result (p=0.05) would lie relative to the background distribution.

**Table S4.**
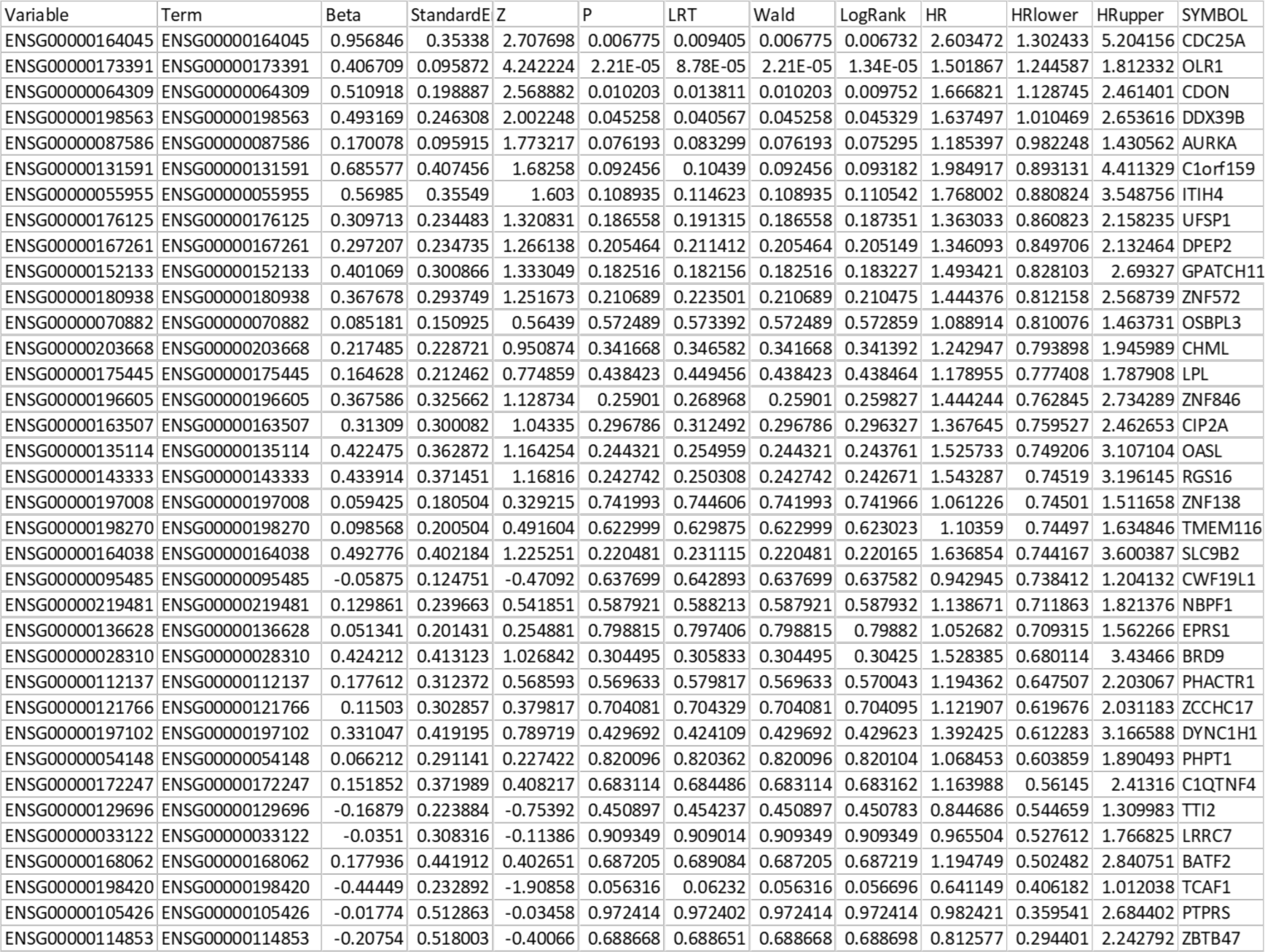
Cox proportional hazard analysis of genes independently associated with biochemical progression-free survival.

**Figure S11.**
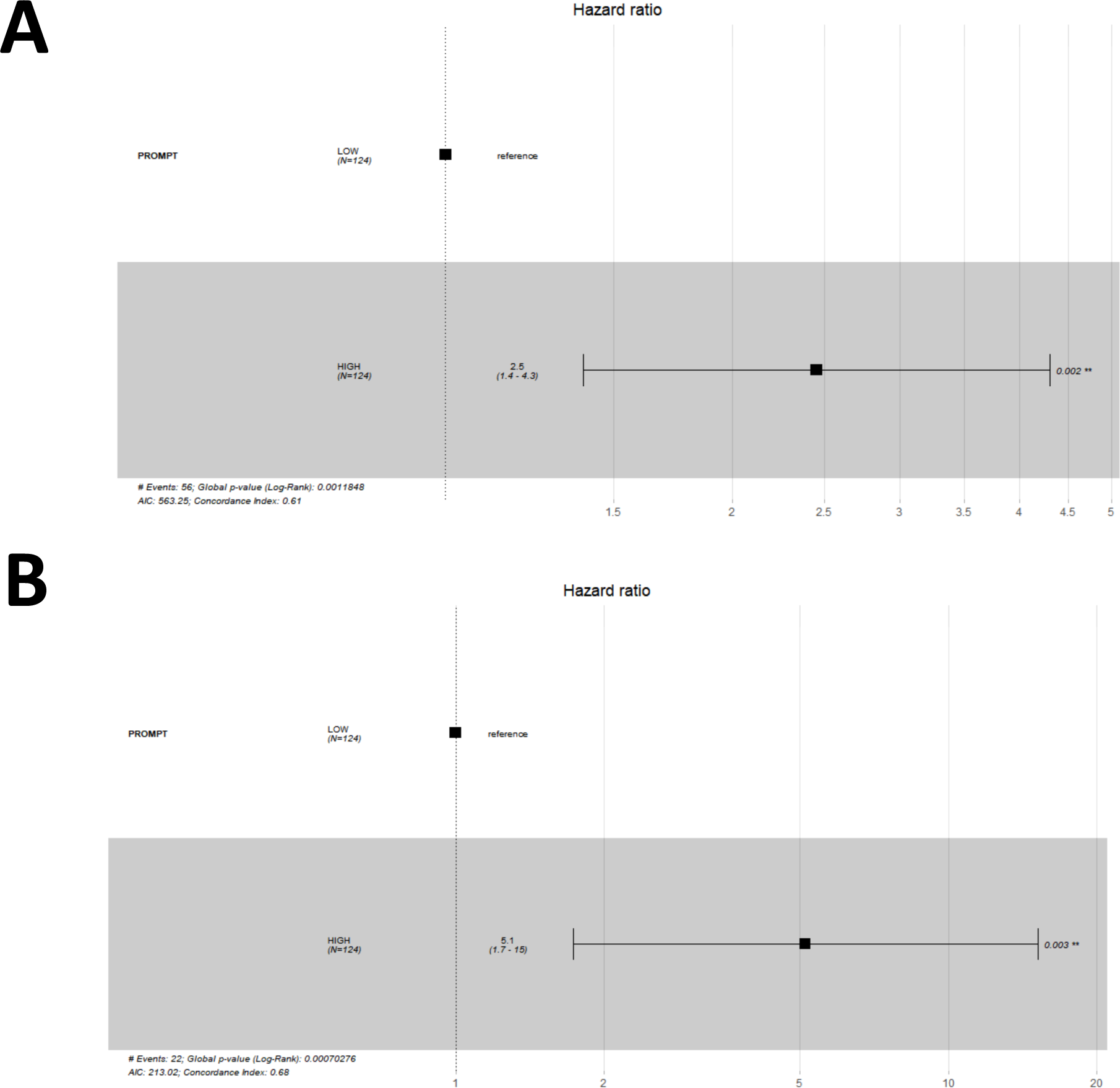
37-gene receiver operator curve for biochemical progression-free survival in the Jain radiotherapy dataset (A). 37-gene receiver operator curve for biochemical progression-free survival in the Jain radiotherapy dataset (B).

**Figure S12.**
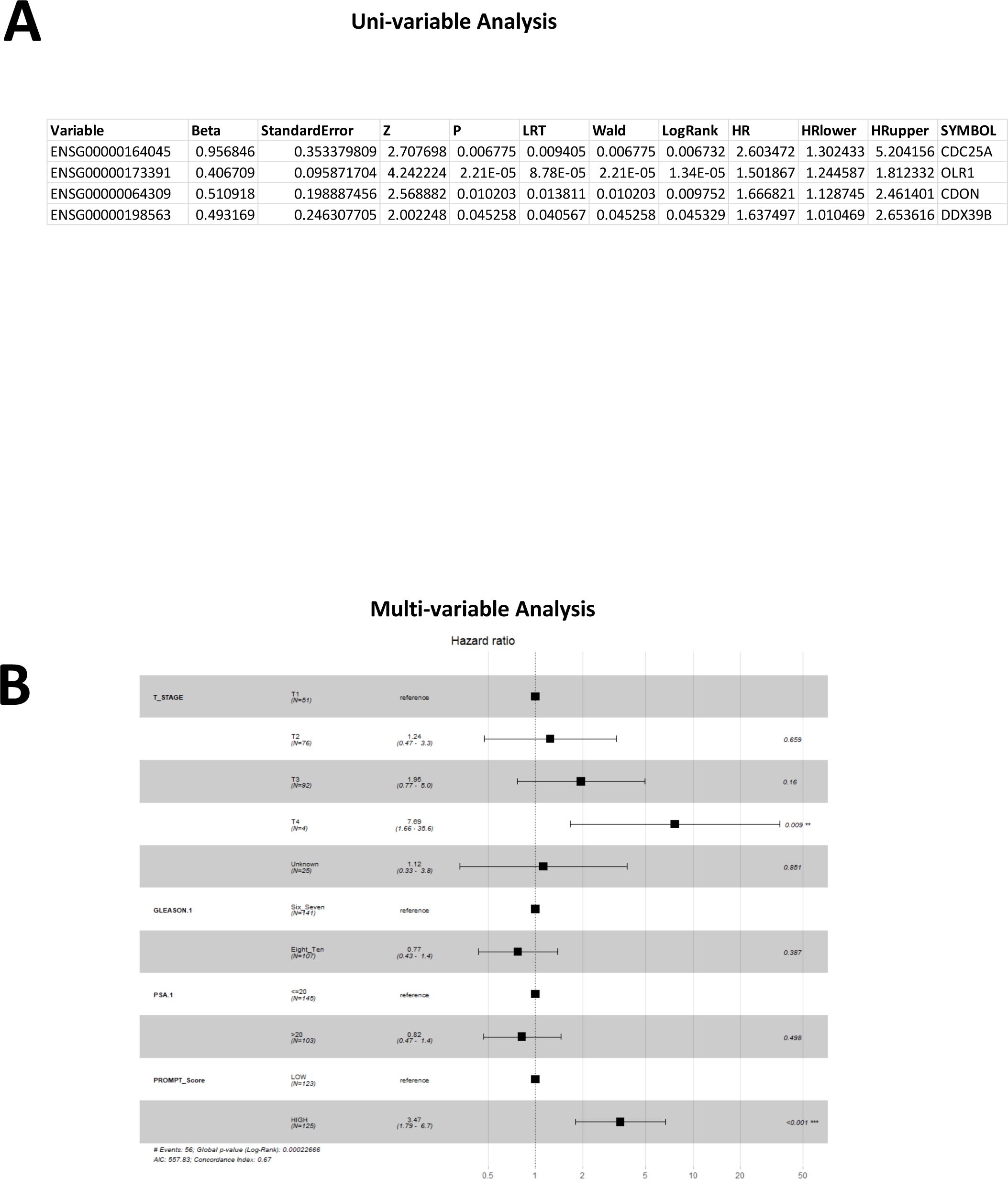
Univariable analysis identifies 4 genes having a lower HR value >1 in the Jain Radiotherapy Cohort.4-gene (mean per gene - median cut-off for cohort) **(A)**. A Cox proportional hazards model analysis of biochemical progression-free survival was performed **(B)**. Forest plot of the cox proportional hazard model for biochemical progression-free survival, incorporating clinicopathological features (T-stage, Gleason score and PSA using clinically relevant cut-points) and high/low expression cohorts (mean per gene - median value threshold for dividing cohort) of the 4 gene signature in the Jain radiotherapy cohort (B).

**Figure S13.**
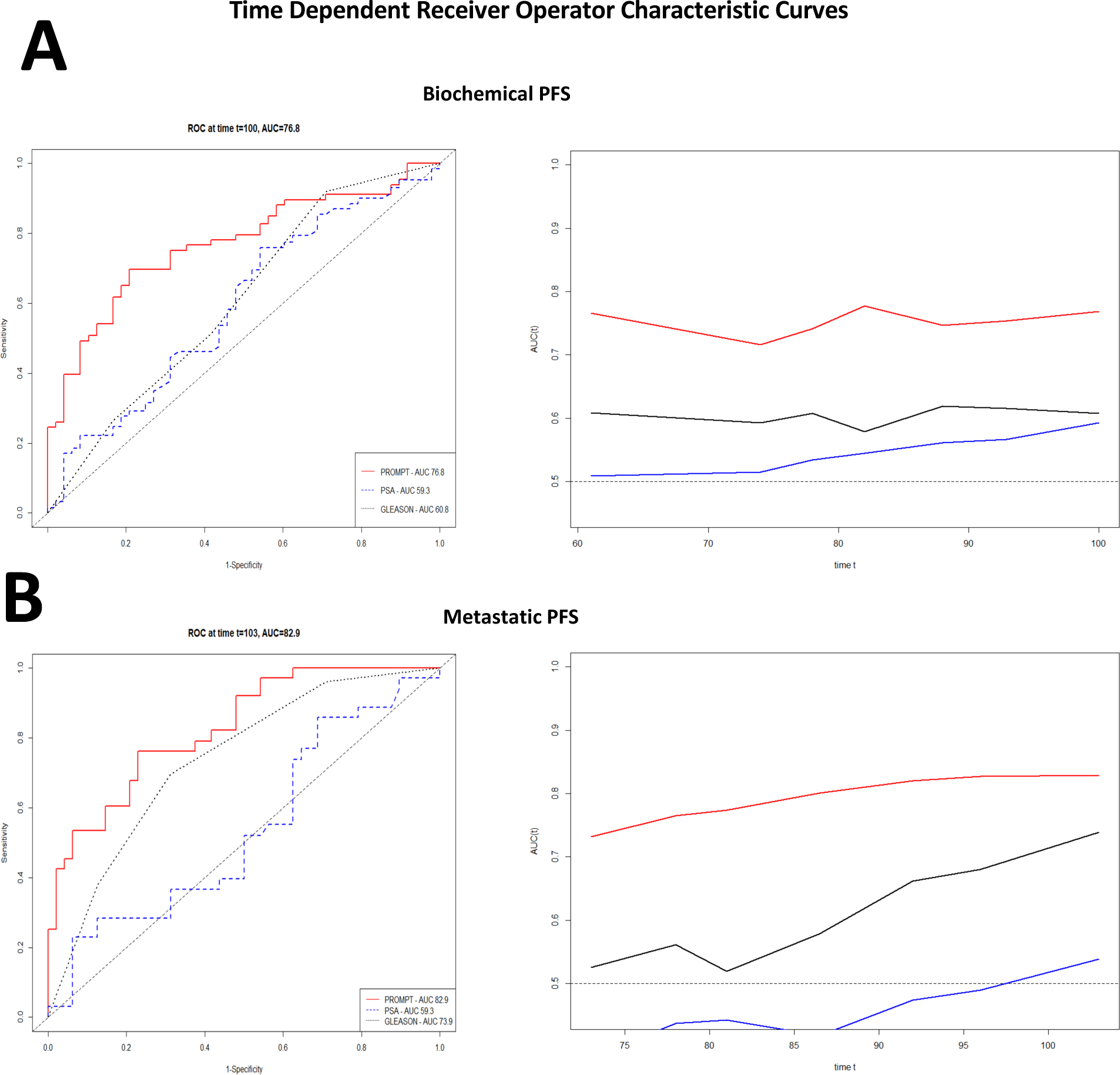
Time dependent Receiver Operator Characteristic curves for biochemical (A) and metastatic (B) progression-free survival in the Jain radiotherapy dataset, for the 4 gene signature, PSA and Gleason sum score.

**Table S5.**
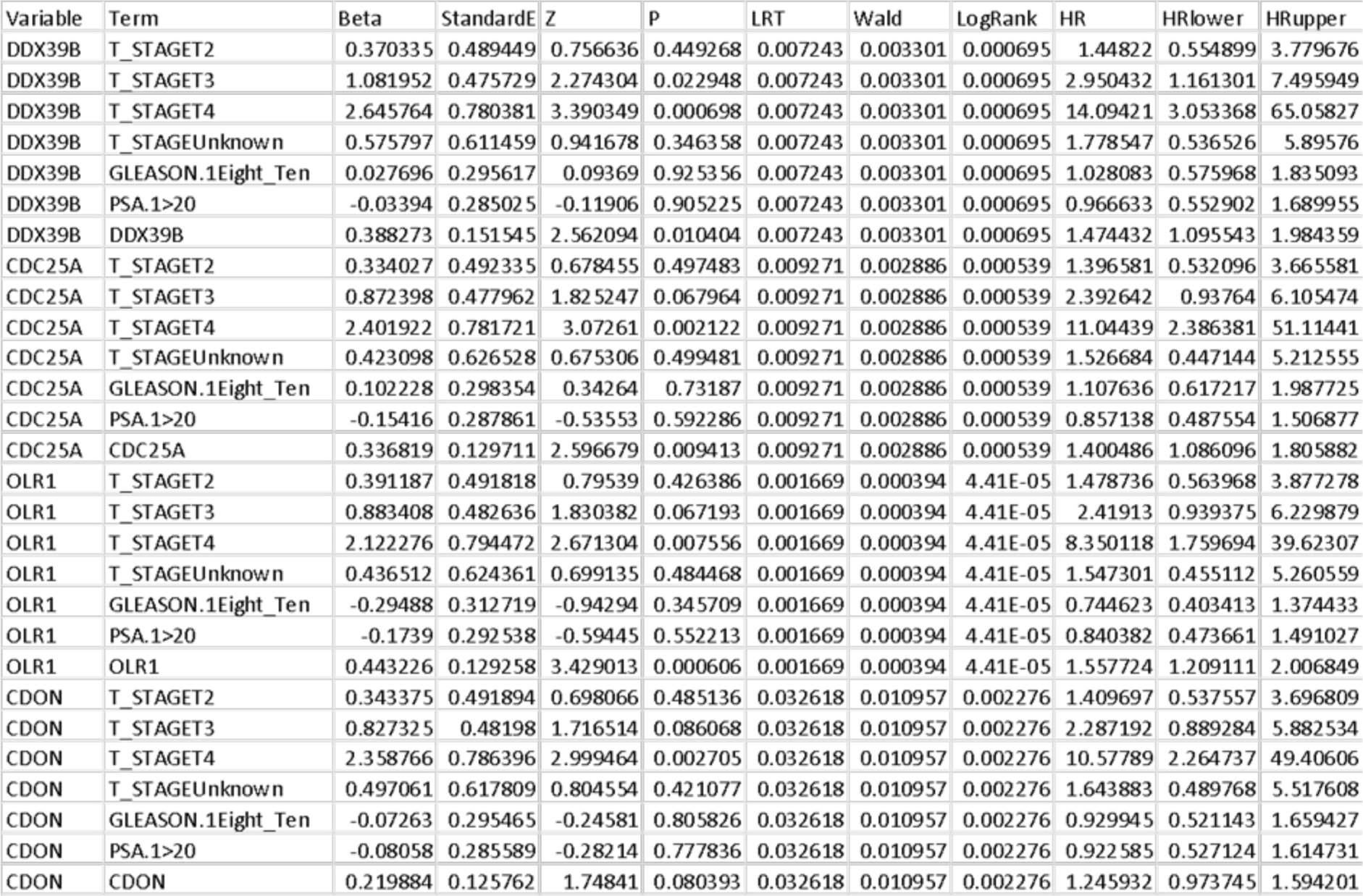
4 gene (mean per gene - median cut-off for cohort) cox proportional hazards and clinical variables.

**Figure S14.**
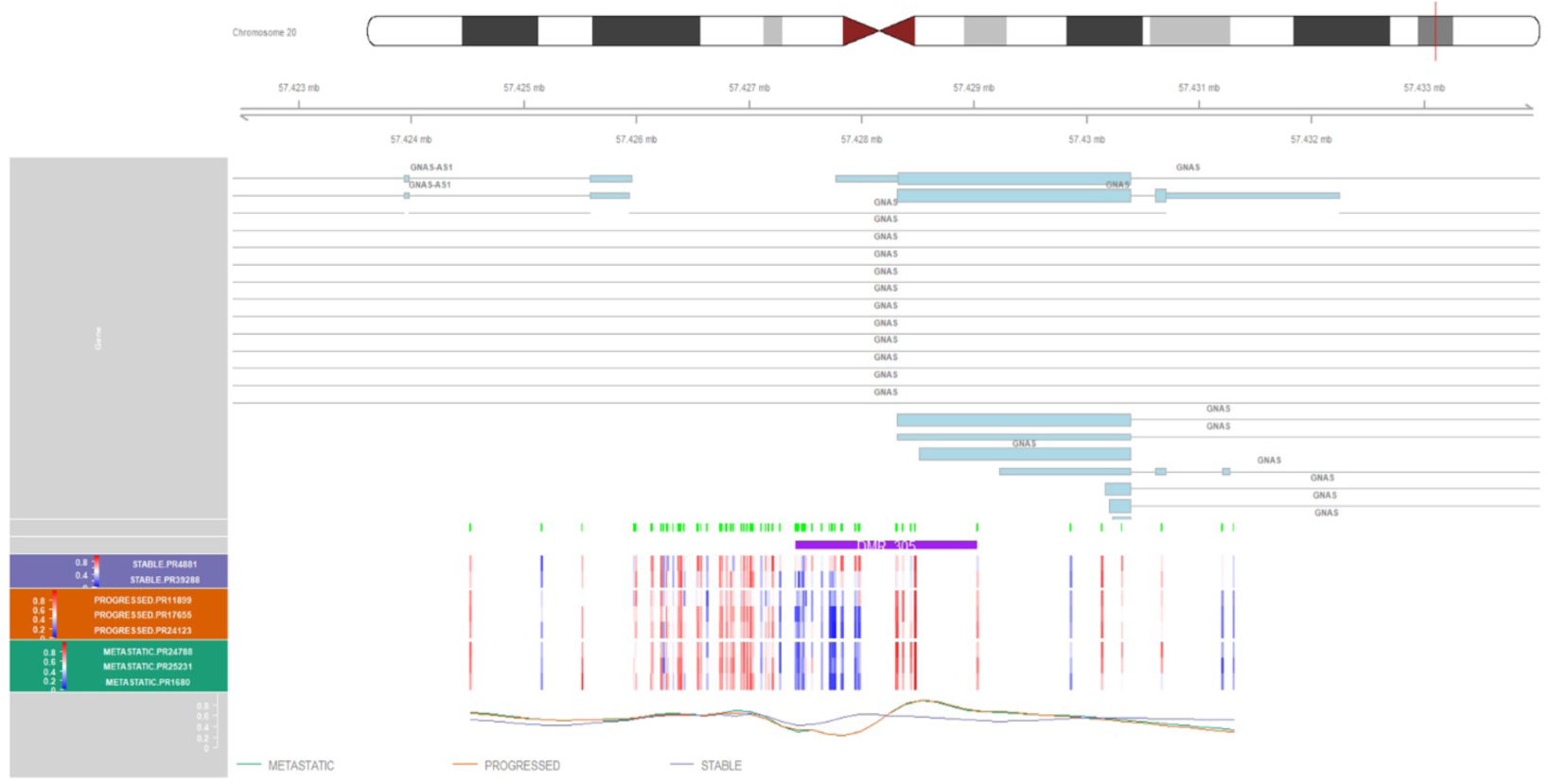
Differentially Methylated Region plot for the *GNAS* gene, visualizing differential methylation between progressed and metastatic cases versus stable cases post-RRT. Genomic co-ordinates and proximal coding regions (top), and heatmap and mean methylation plots (bottom), are illustrated.

**Figure S15.**
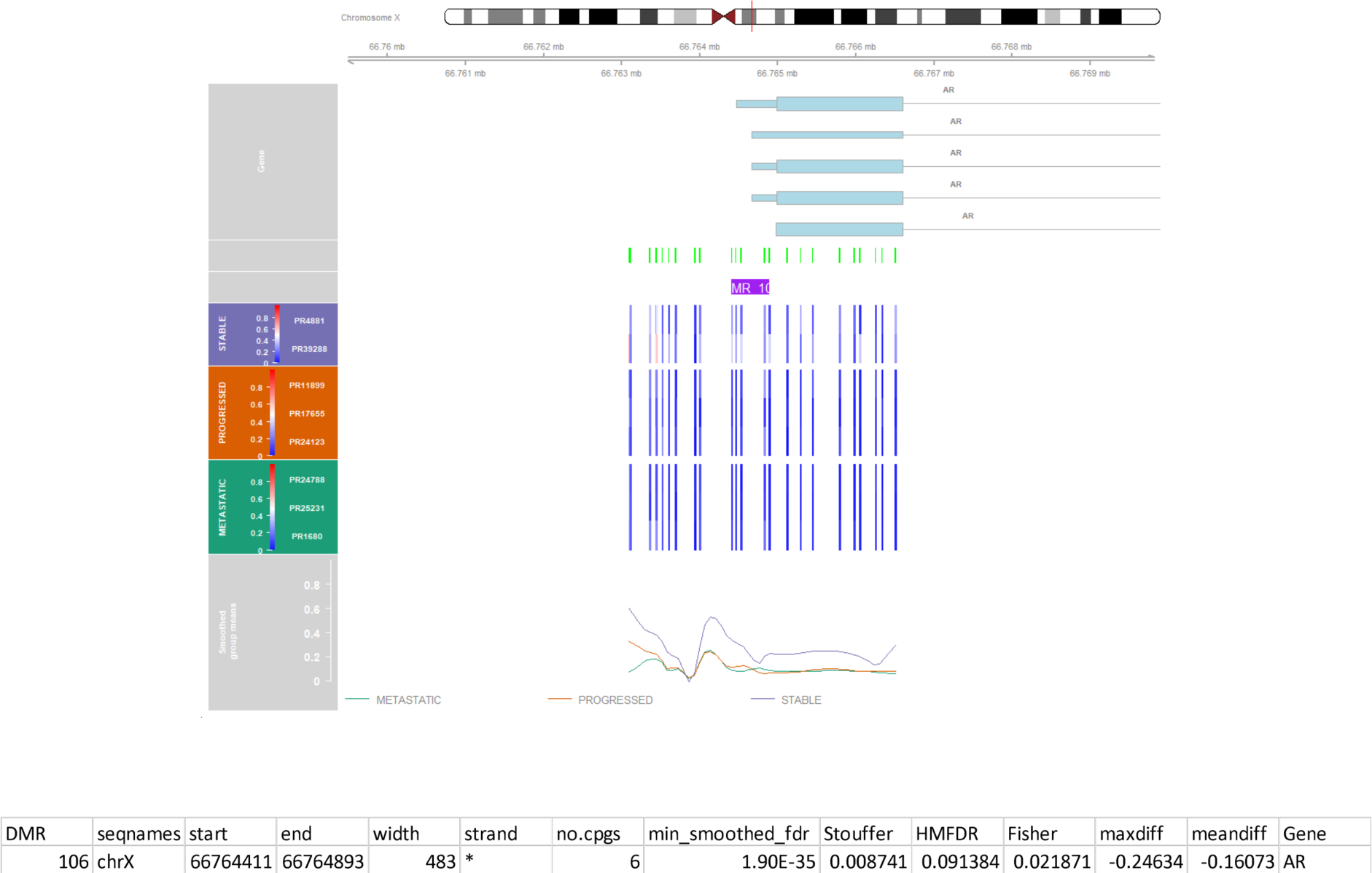
Differentially Methylated Region plot for the *Androgen Receptor* gene, visualizing differential methylation between progressed and metastatic cases versus stable cases post-RRT. Genomic co-ordinates and proximal coding regions (top), and heatmap and mean methylation plots (bottom), are illustrated.

**Figure S16.**
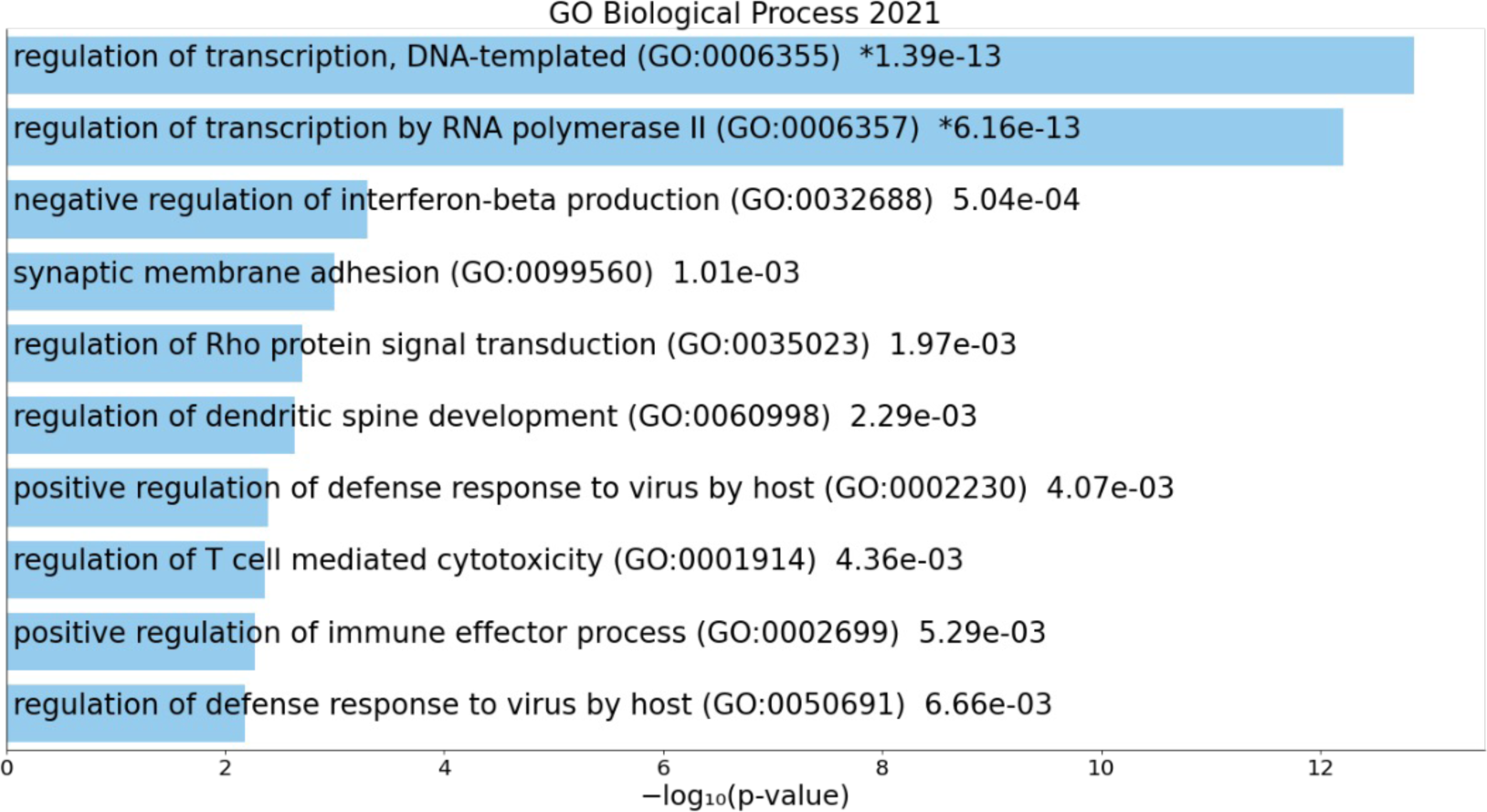
Bar-plot of over-representation analysis of Chromosome 19 (enrichr) GO Biological Process pathways, and Differentially Methylated Genes, between progressed and metastatic cases versus stable cases.

**Figure S17.**
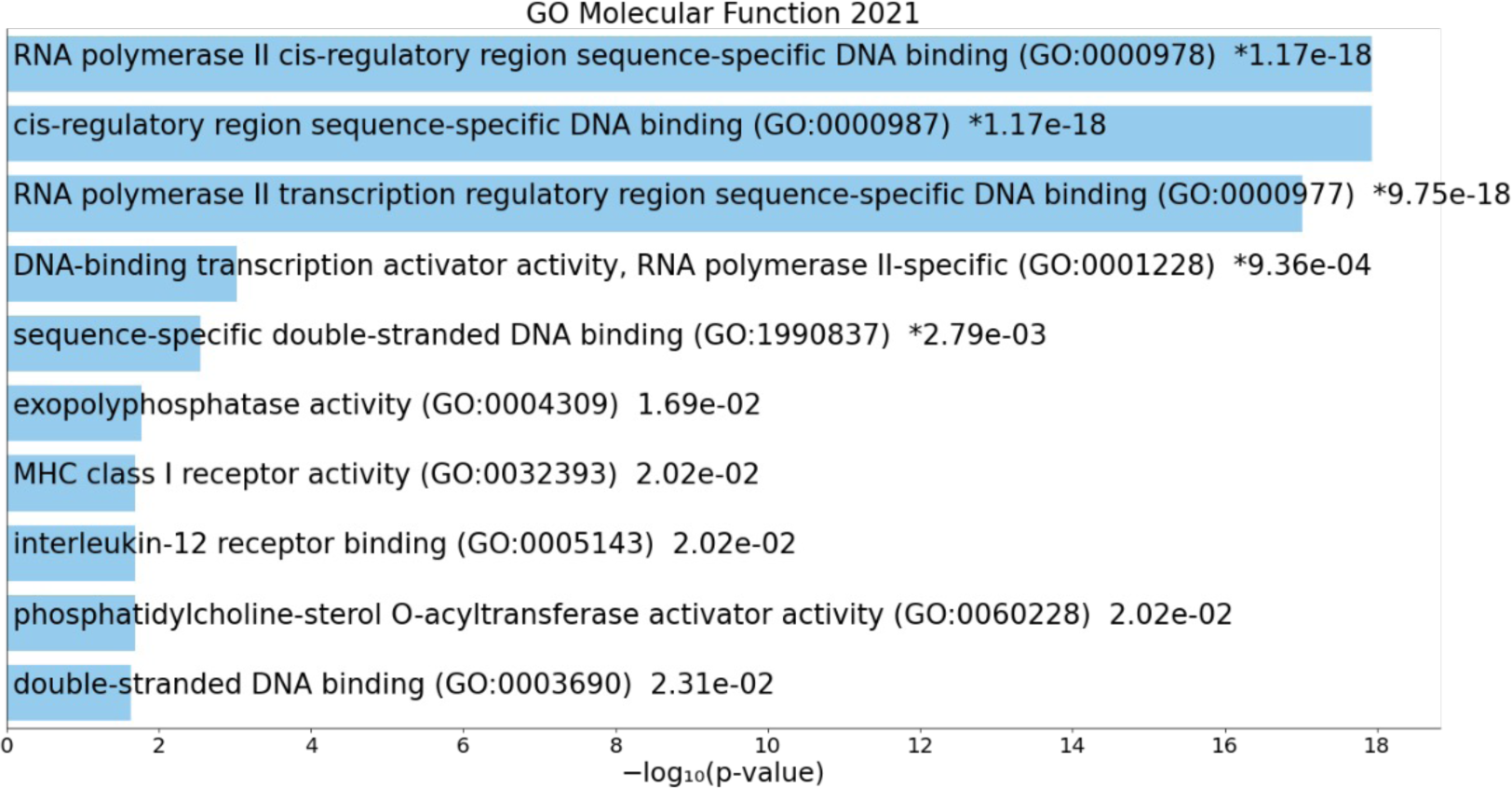
Bar-plot of over-representation analysis of Chromosome 19 (enrichr) GO Molecular Function pathways, and Differentially Methylated Genes, between progressed and metastatic cases versus stable cases.

**Figure S18.**
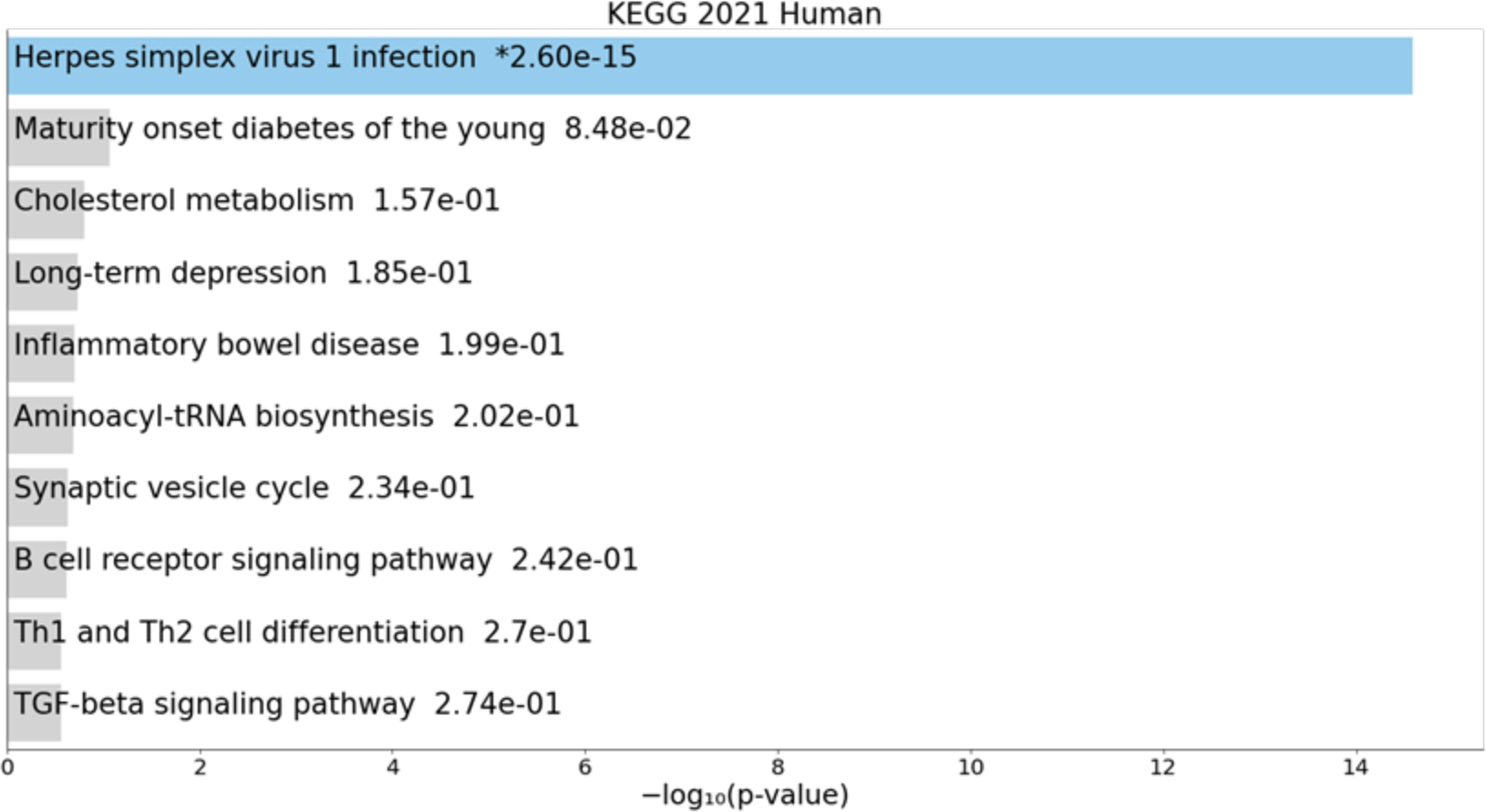
Bar-plot of over-representation analysis of Chromosome 19 (enrichr) KEGG pathways, and Differentially Methylated Genes, between progressed and metastatic cases versus stable cases.

**Figure S19.**
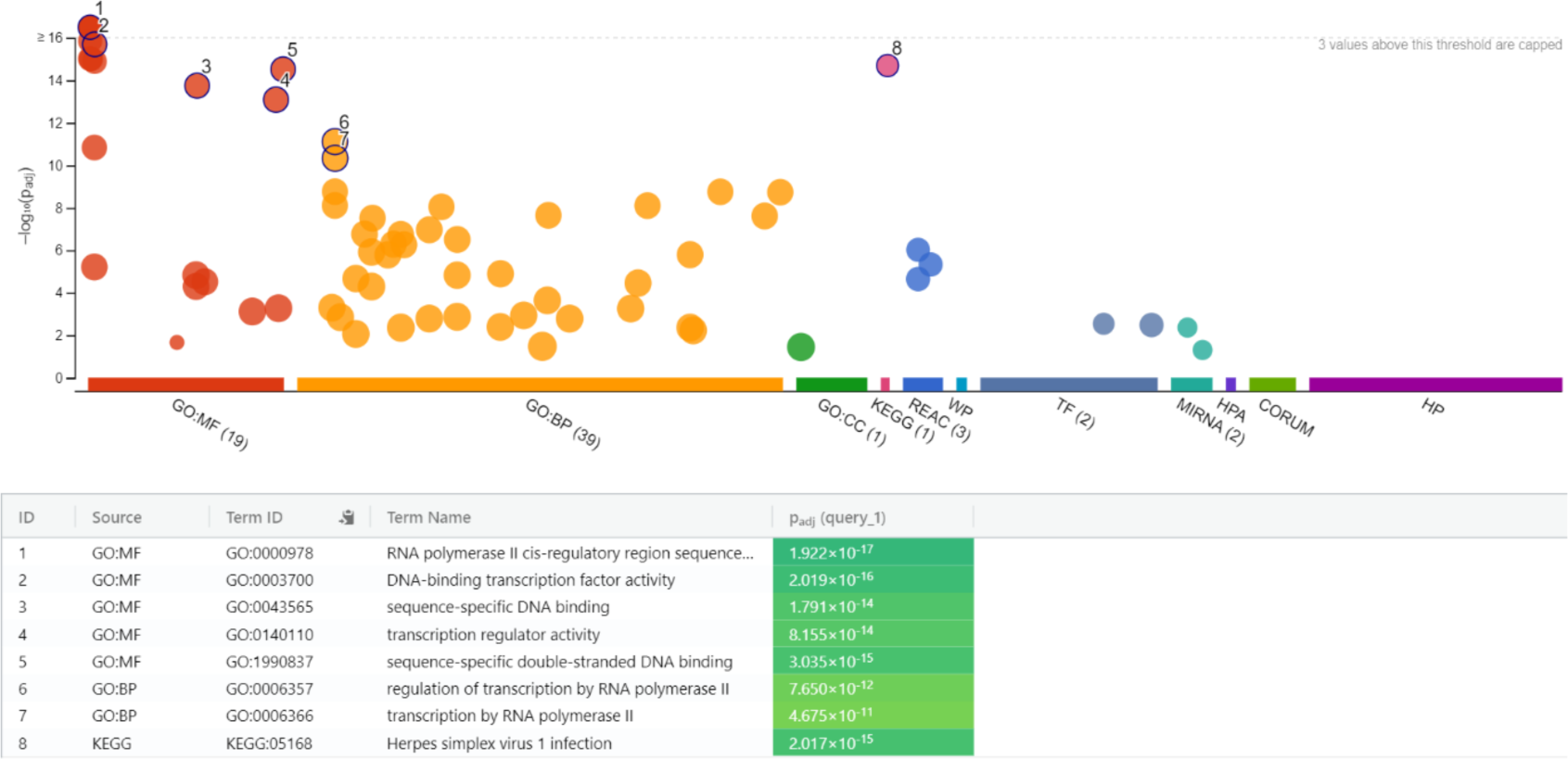
Over-representation analysis of all pathways for Chromosome 19 (gprofiler).

**Figure S20.**
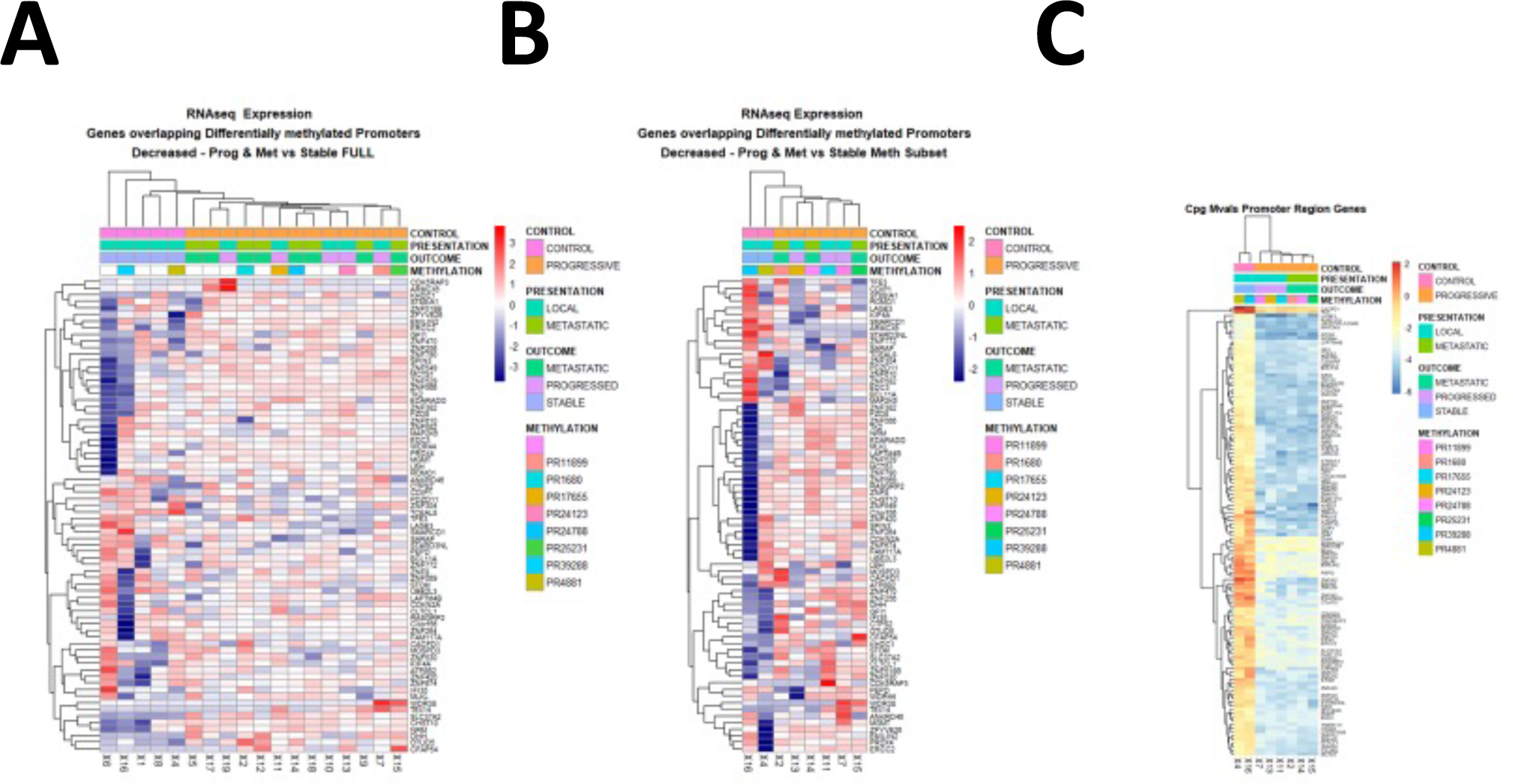
Heatmap analysis of RNA expression of differentially hypo-methylated genes at promoter in the full cohort, methylation subset, and cpg Mvals for methylation. Quantseq RNA expression analysis heatmap of genes with differentially hypo-methylated promoter regions between progressed and metastatic cases versus stable cases in the whole cohort. (**A**). Quantseq RNA expression analysis heatmap of genes with differentially hypo-methylated promoter regions between progressed and metastatic cases versus stable cases in the methylation analysis cohort (**B**). Methylation (Mvals) heatmap of Cpgs in promoter regions differentially methylated between progressed and metastatic cases versus stable cases (**C**).

**Figure S21.**
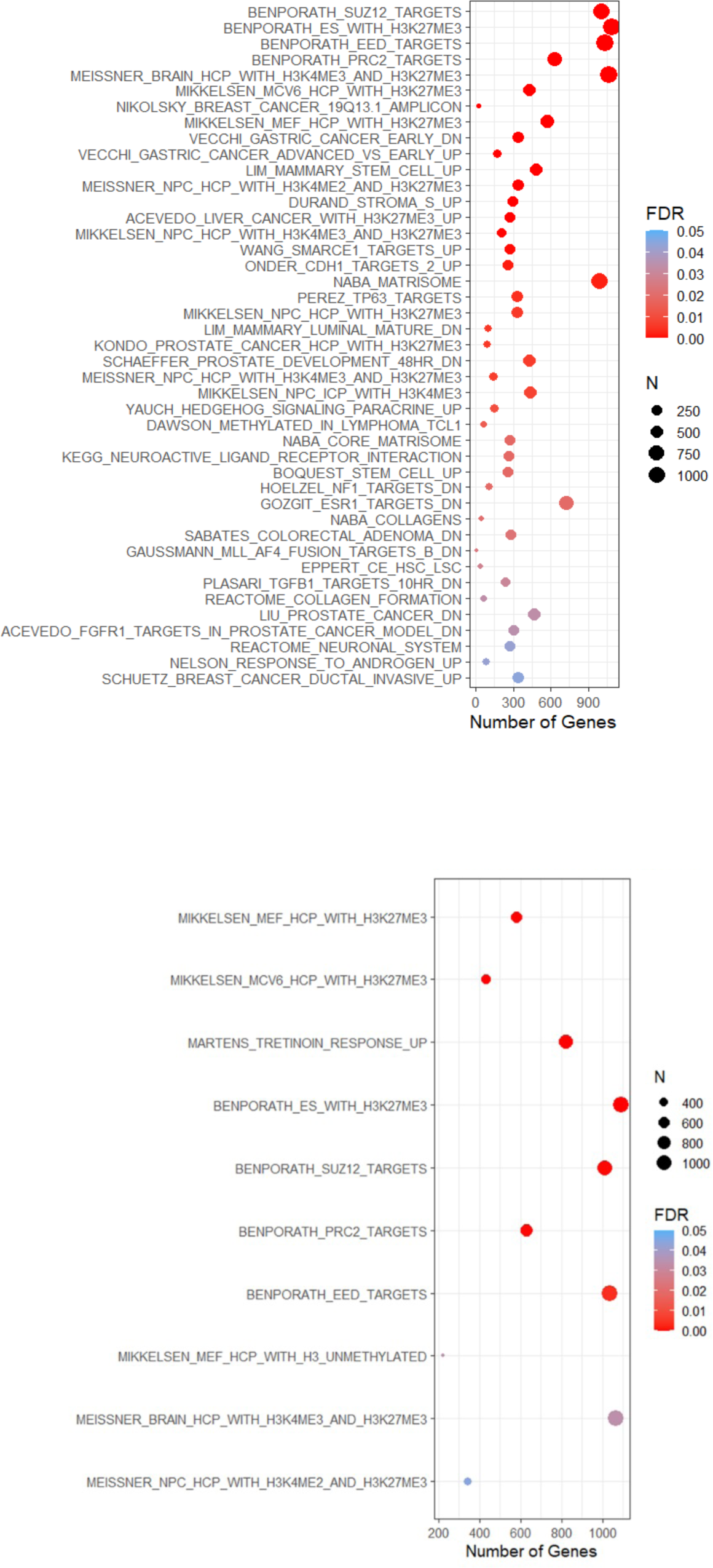
Dot-plot over-representation analysis of hypomethylated (**A**) and hypermethylated (**B**) C2 Curated Pathway (MSigDB) genes in progressed and metastatic cases versus stable cases post-RT.

**Figure S22.**
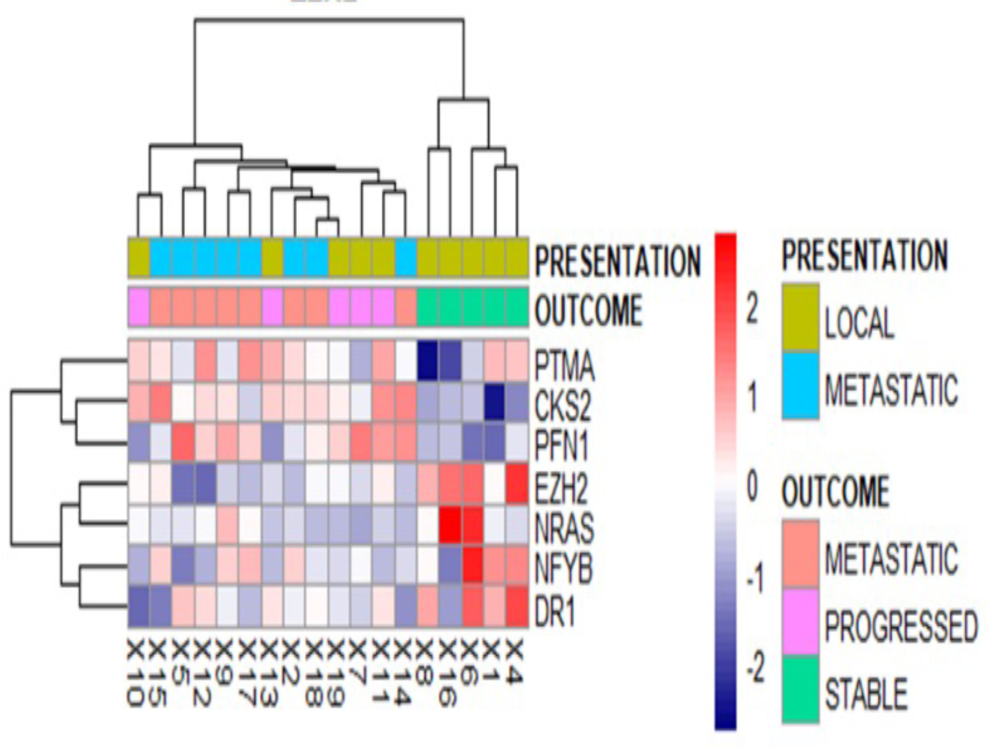
Heatmap of EZH2 expression, and expression of EZH2 target genes, from the Quantseq data, demonstrating significant differential expression between progressed and metastatic cases versus stable cases (padj<0.05).

**Figure S23.**
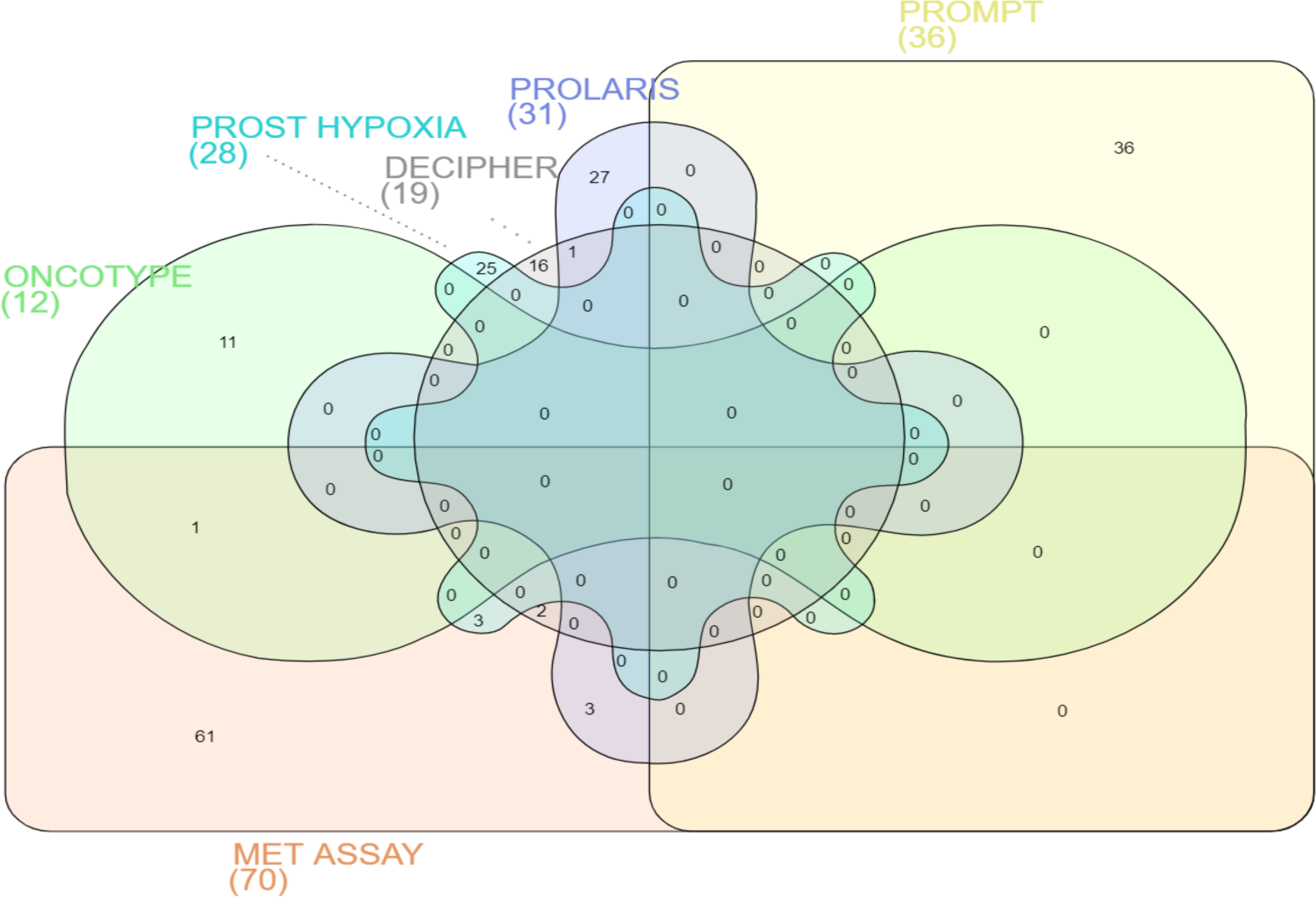
Signature Genelist overlap.

## Supplementary Methods

### Gene Expression & Methylation Analysis

3’RNAseq FastQ files were concatenated for each patient, and polyA and llumina adapter trimming (AGATCGGAAGAGC) was performed using trimmomatic (v0.25), prior to alignment with STAR (2.7.7a) to Hg38 (GRCh38.95), with featurecounts used to generate counts exact parameters are available upon request. Filtering of lowly expressed genes was performed, followed by differential expression analysis using DESEQ2 (v1.26.0)^22^. Over-representation analysis was performed using Clusterprofiler, enrichr and gprofiler. The gene set variation analysis GSVA (v1.34.0) package was used to perform single sample gene set enrichment analysis (ssGSEA)^23^. C2 Curated genesets (c2.all.v7.2.entrez.gmt) were used from Molecular Signatures Database (MSigDB) website (https://www.gsea-msigdb.org/gsea/msigdb).

For DNA methylation analysis, QC, filtering of poor performing probes, cross-reactive probes and normalization was performed prior to differential methylation analysis using minfi and MissMethyl package (v1.32.0). The manifest file “infinium.methylationepic.v.1.0.b5” was used. DMRcate package (2.0.7) was used for differential methylation and Granges for visualization, MissMethyl (v1.32.0) was used for pathway analyses.

### Additional Analysis

Sequencing and clinical data from the external dataset GSE116918 were downloaded from Gene Expression Omnibus (GEO available at https://www.ncbi.nlm.nih.gov/geo/). Probes were filtered to only include non-overlapping exonic probes, and multiple probes were merged to the mean value per gene. To overcome technology platform differences, as GSE116918 utilized a different sequencing technology (microarray) to those used in this study 3’RNAseq, mean values of fully exonic probes for the relevant genes were used from the GSE116918 dataset. NanoString probes are designed to span exons, such that only mature mRNAs are counted. The Quantseq method sequences mature transcripts with polyA tails, and this was used with the FeatureCounts and Subread package for quantification and identifications of alternate transcripts. Survminer and Survival packages were used for survival analysis with cox proportional hazards models utilizing known clinicopathological features, staging and Gleason score. Survival analysis was performed using a Cox proportional hazards model comprising clinicopathological features and expression of selected genes, followed by estimation of time-dependent Receiver Operator Characteristics (ROC) using TimeROC (package 0.4). KMunicate (0.2.0) and survival (3.2-3) packages were used for survival analysis, with extended risk tables for time to event analysis.

R sessionInfo() is provided in the Package Summary file.

