## Supplementary Package Summary for "Molecular analysis of archival diagnostic prostate cancer biopsies identifies genomic similarities in cases with progression post-radiotherapy, and those with *de novo* metastatic disease"

R version 3.6.2 (2019-12-12)

Platform: x86_64-w64-mingw32/x64 (64-bit)

Running under: Windows 10 x64 (build 19045)

Matrix products: default

locale:

[1] LC_COLLATE=English_United Kingdom.1252 LC_CTYPE=English_United Kingdom.1252

[3] LC_MONETARY=English_United Kingdom.1252 LC_NUMERIC=C

[5] LC_TIME=English_United Kingdom.1252

attached base packages:

[1] grid	stats4	parallel stats	graphics grDevices utils	datasets

[9] methods	base

other attached packages:

[1] ReactomePA_1.30.0

[2] edgeR_3.28.1

[3] EBSeqHMM_1.20.0

[4] EBSeq_1.26.0

[5] testthat_2.3.2

[6] gplots_3.0.4

[7] blockmodeling_1.0.0

[8] ComplexHeatmap_2.2.0

[9] circlize_0.4.10

[10] DMRcate_2.0.7

[11] Gviz_1.30.3

[12] IlluminaHumanMethylation450kanno.ilmn12.hg19_0.6.0

[13] knitr_1.29

[14] ruv_0.9.7.1

[15] statmod_1.4.34

[16] shinyMethyl_1.22.0

[17] IlluminaHumanMethylation450kmanifest_0.4.0

[18] minfi_1.32.0

[19] bumphunter_1.28.0

[20] locfit_1.5-9.4

[21] iterators_1.0.12

[22] foreach_1.5.0

[23] Biostrings_2.54.0

[24] XVector_0.26.0

[25] shiny_1.5.0

[26] VennDiagram_1.6.20

[27] futile.logger_1.4.3

[28] missMethyl_1.20.4

[29] DESeq2_1.26.0

[30] SummarizedExperiment_1.16.1

[31] DelayedArray_0.12.3

[32] matrixStats_0.56.0

[33] GenomicRanges_1.38.0

[34] GenomeInfoDb_1.22.1

[35] GSVA_1.34.0

[36] enrichplot_1.6.1

[37] clusterProfiler_3.14.3

[38] cowplot_1.0.0

[39] KMunicate_0.2.0

[40] org.Hs.eg.db_3.10.0

[41] stringr_1.4.0

[42] dplyr_1.0.0

[43] gridExtra_2.3

[44] survminer_0.4.8

[45] ggpubr_0.4.0

[46] ggplot2_3.3.2

[47] RColorBrewer_1.1-2

[48] pheatmap_1.0.12

[49] GEOquery_2.54.1

[50] limma_3.42.2

[51] SigCheck_2.18.0

[52] survival_3.2-3

[53] BiocParallel_1.20.1

[54] e1071_1.7-4

[55] MLInterfaces_1.66.5

[56] cluster_2.1.0

[57] GSEABase_1.48.0

[58] graph_1.64.0

[59] annotate_1.64.0

[60] XML_3.99-0.3

[61] AnnotationDbi_1.48.0

[62] IRanges_2.20.2

[63] S4Vectors_0.24.4

[64] Biobase_2.46.0

[65] BiocGenerics_0.32.0

loaded via a namespace (and not attached):

[1] tinytex_0.25

[2] graphlayouts_0.7.0

[3] lattice_0.20-41

[4] haven_2.3.1

[5] graphite_1.32.0

[6] vctrs_0.3.1

[7] mgcv_1.8-33

[8] beanplot_1.2

[9] DSS_2.34.0

[10] blob_1.2.1

[11] prodlim_2019.11.13

[12] later_1.1.0.1

[13] nloptr_1.2.2.2

[14] DBI_1.1.0

[15] R.utils_2.10.1

[16] rappdirs_0.3.1

[17] jpeg_0.1-8.1

[18] zlibbioc_1.32.0

[19] MatrixModels_0.4-1

[20] pls_2.8-0

[21] htmlwidgets_1.5.3

[22] mvtnorm_1.1-1

[23] GlobalOptions_0.1.2

[24] DEoptimR_1.0-9

[25] illuminaio_0.28.0

[26] tidygraph_1.2.0

[27] Rcpp_1.0.5

[28] readr_1.3.1

[29] KernSmooth_2.23-17

[30] promises_1.1.1

[31] gdata_2.18.0

[32] methylumi_2.32.0

[33] Hmisc_4.4-1

[34] RSpectra_0.16-0

[35] fastmatch_1.1-0

[36] digest_0.6.25

[37] png_0.1-7

[38] polspline_1.1.19

[39] nor1mix_1.3-0

[40] DOSE_3.12.0

[41] ggraph_2.0.3

[42] pkgconfig_2.0.3

[43] GO.db_3.10.0

[44] DelayedMatrixStats_1.8.0

[45] minqa_1.2.4

[46] reticulate_1.18

[47] modeltools_0.2-23

[48] GetoptLong_1.0.4

[49] xfun_0.15

[50] zoo_1.8-8

[51] tidyselect_1.1.0

[52] reshape2_1.4.4

[53] purrr_0.3.4

[54] kernlab_0.9-29

[55] viridisLite_0.3.0

[56] rtracklayer_1.46.0

[57] rlang_0.4.6

[58] glue_1.4.1

[59] ensembldb_2.10.2

[60] fpc_2.2-9

[61] lambda.r_1.2.4

[62] umap_0.2.7.0

[63] lava_1.6.8

[64] europepmc_0.4

[65] ggsignif_0.6.0

[66] threejs_0.3.3

[67] SparseM_1.78

[68] httpuv_1.5.4

[69] class_7.3-17

[70] preprocessCore_1.48.0

[71] TH.data_1.0-10

[72] reactome.db_1.70.0

[73] DO.db_2.9

[74] bsseq_1.22.0

[75] jsonlite_1.7.0

[76] bit_4.0.4

[77] mime_0.9

[78] Rsamtools_2.2.3

[79] stringi_1.4.6

[80] pammtools_0.5.6

[81] quadprog_1.5-8

[82] TcGSA_0.12.7

[83] bitops_1.0-6

[84] RSQLite_2.2.0

[85] tidyr_1.1.0

[86] data.table_1.13.0

[87] rstudioapi_0.11

[88] GenomicAlignments_1.22.1

[89] sfsmisc_1.1-11

[90] nlme_3.1-149

[91] qvalue_2.18.0

[92] gbm_2.1.8

[93] VariantAnnotation_1.32.0

[94] gridGraphics_0.5-0

[95] survMisc_0.5.5

[96] R.oo_1.24.0

[97] prabclus_2.3-2

[98] dbplyr_1.4.4

[99] readxl_1.3.1

[100] lifecycle_0.2.0

[101] ExperimentHub_1.12.0

[102] munsell_0.5.0

[103] cellranger_1.1.0

[104] R.methodsS3_1.8.1

[105] hwriter_1.3.2

[106] caTools_1.18.0

[107] codetools_0.2-16

[108] htmlTable_2.0.1

[109] triebeard_0.3.0

[110] xtable_1.8-4

[111] diptest_0.76-0

[112] formatR_1.7

[113] BiocManager_1.30.10

[114] abind_1.4-5

[115] farver_2.0.3

[116] km.ci_0.5-2

[117] AnnotationHub_2.18.0

[118] ggvis_0.4.7

[119] askpass_1.1

[120] biovizBase_1.34.1

[121] IlluminaHumanMethylationEPICanno.ilm10b4.hg19_0.6.0

[122] shinythemes_1.1.2

[123] tibble_3.0.2

[124] futile.options_1.0.1

[125] dichromat_2.0-0

[126] Matrix_1.2-18

[127] ellipsis_0.3.1

[128] prettyunits_1.1.1

[129] ggridges_0.5.3

[130] mclust_5.4.6

[131] igraph_1.2.5

[132] multtest_2.42.0

[133] mlbench_2.1-3

[134] fgsea_1.12.0

[135] pec_2019.11.03

[136] htmltools_0.5.1.1

[137] BiocFileCache_1.10.2

[138] yaml_2.2.1

[139] GenomicFeatures_1.38.2

[140] interactiveDisplayBase_1.24.0

[141] foreign_0.8-72

[142] withr_2.2.0

[143] bit64_4.0.5

[144] BiasedUrn_1.07

[145] rngtools_1.5

[146] doRNG_1.8.2

[147] multcomp_1.4-14

[148] robustbase_0.93-6

[149] ProtGenerics_1.18.0

[150] GSA_1.03.1

[151] GOSemSim_2.12.1

[152] memoise_1.1.0

[153] forcats_0.5.0

[154] rio_0.5.16

[155] geneplotter_1.64.0

[156] permute_0.9-5

[157] curl_4.3

[158] urltools_1.7.3

[159] conquer_1.0.2

[160] checkmate_2.0.0

[161] rjson_0.2.20

[162] openxlsx_4.2.2

[163] rstatix_0.6.0

[164] ggrepel_0.8.2

[165] clue_0.3-57

[166] tools_3.6.2

[167] sandwich_3.0-0

[168] magrittr_1.5

[169] RCurl_1.98-1.2

[170] timeROC_0.4

[171] car_3.0-9

[172] ggplotify_0.0.5

[173] xml2_1.3.2

[174] httr_1.4.2

[175] assertthat_0.2.1

[176] boot_1.3-25

[177] R6_2.4.1

[178] Rhdf5lib_1.8.0

[179] AnnotationFilter_1.10.0

[180] nnet_7.3-14

[181] progress_1.2.2

[182] genefilter_1.68.0

[183] gtools_3.8.2

[184] shape_1.4.5

[185] BiocVersion_3.10.1

[186] HDF5Array_1.14.4

[187] rhdf5_2.30.1

[188] splines_3.6.2

[189] carData_3.0-4

[190] colorspace_1.4-1

[191] generics_0.1.1

[192] base64enc_0.1-3

[193] pillar_1.4.6

[194] IlluminaHumanMethylationEPICmanifest_0.3.0

[195] tweenr_1.0.1

[196] rvcheck_0.1.8

[197] GenomeInfoDbData_1.2.2

[198] plyr_1.8.6

[199] gtable_0.3.0

[200] timereg_1.9.8

[201] zip_2.1.1

[202] latticeExtra_0.6-29

[203] biomaRt_2.42.1

[204] fastmap_1.0.1

[205] crosstalk_1.1.0.1

[206] pscl_1.5.5

[207] flexmix_2.3-17

[208] quantreg_5.67

[209] broom_0.7.0

[210] openssl_1.4.2

[211] BSgenome_1.54.0

[212] scales_1.1.1

[213] backports_1.1.7

[214] base64_2.0

[215] rms_6.0-1

[216] lme4_1.1-23

[217] hms_0.5.3

[218] ggforce_0.3.2

[219] scrime_1.3.5

[220] KMsurv_0.1-5

[221] polyclip_1.10-0

[222] numDeriv_2016.8-1.1

[223] siggenes_1.60.0

[224] lazyeval_0.2.2

[225] Formula_1.2-3

[226] crayon_1.3.4

[227] MASS_7.3-52

[228] viridis_0.5.1

[229] reshape_0.8.8

[230] rpart_4.1-15

[231] compiler_3.6.2

>
